# Robustness of Hill’s overlapping-generation method for calculating *N_e_* to extreme patterns of reproductive success

**DOI:** 10.1101/2023.02.13.528382

**Authors:** Robin S. Waples

## Abstract

For species with overlapping generations, the most widely-used method to calculate effective population size (*N_e_*) is Hill’s, the key parameter for which is lifetime variance in offspring number (*V*_*k*•_). Hill’s model assumes stable age structure and constant abundance, and sensitivity to those assumptions has been evaluated previously. Here I evaluate robustness of Hill’s model to extreme patterns of reproductive success, whose effects have not been previously examined: 1) very strong reproductive skew; 2) strong temporal autocorrelations in individual reproductive success; and 3) strong covariance of individual reproduction and survival. Genetic drift (loss of heterozygosity and increase in allele-frequency variance) was simulated in age-structured populations using methods that: generated no autocorrelations or covariances (Model NoCor); or created strong positive (Model Positive) or strong negative (Model Negative) temporal autocorrelations in reproduction and covariances between reproduction and survival. Compared to Model NoCor, the other models led to greatly elevated or reduced *V*_*k*•_, and hence greatly reduced or elevated *N_e_*, respectively. A new index is introduced (ρ*_α_*,*_α_***_+_**), which is the correlation between 1) the number of offspring produced by each individual at the age at maturity (α), and 2) the total number of offspring produced during the rest of their lifetimes. Mean ρ*_α_*,*_α_***_+_** was ≈0 under Model NoCor, strongly positive under Model Positive, and strongly negative under Model Negative. Even under the most extreme reproductive scenarios in Models Positive and Negative, when *V*_*k*•_ was calculated from the realized population pedigree and used to calculate *N_e_* in Hill’s model, the result accurately predicted the rate of genetic drift in simulated populations. These results held for scenarios where age-specific reproductive skew was random (variance≈mean) and highly overdispersed (variance up to 20 times the mean). Collectively, these results are good news for researchers as they demonstrate the robustness of Hill’s model even to extreme repro0ductive scenarios.

## INTRODUCTION

The majority of populations in nature are age structured, but most evolutionary theory was originally developed for models that assume discrete generations. Considerable progress has been made in adapting discrete-generation theory to account for age structure (e.g., Charlesworth 1994; Cushing et al. 1994; Lande et al. 2003), but this process remains challenging.

One of the most important parameters in evolutionary biology is effective population size (*N_e_*), which, in addition to determining the rate of genetic drift, modulates the effectiveness of natural selection and hence the rate of evolutionary adaptation. Wright’s original concept of *N_e_* (1931, 1938) assumed that generations were discrete. Of the various methods researchers have proposed for incorporating age structure into the concept of effective size, that of Hill (1972) is the most widely used. Hill showed that, if age-structure is stable and the population produces a constant number (*N_1_*) of offspring in each cohort, *N_e_* per generation is given by

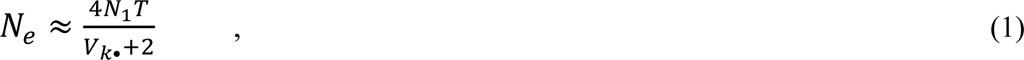

where 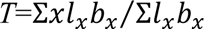 is generation length (average age of parents of the newborn cohort) and *V*_*k*•_ is the variance in lifetime reproductive success (*LRS*), measured as the variance in lifetime number of offspring among the *N_1_* individuals in a cohort.

The assumption in Hill’s model of stable age structure and constant *N* has been extensively evaluated with diverse life histories and found to be robust to random demographic variability (Waples et al. 2011; 2014); these evaluations also found Hill’s model to be robust to skewed adult sex ratios and to modestly overdispersed variance in reproductive success. However, the models used in those evaluations assumed independence of reproduction and survival, which means that the robustness of Hill’s model to temporal autocorrelation in reproduction and covariance of reproduction and survival has not been rigorously tested. This is an important gap, for two major reasons.

First, temporal autocorrelations and covariances are common in many species. The theory of life-history evolution is based on the premise that biological constraints impose intrinsic tradeoffs between reproduction now and reproduction later, and between reproduction and subsequent survival (Williams 1966; Bell 1980; Reznick 1992; Roff 1992). These tradeoffs imply a negative temporal autocorrelation in individual reproduction and a negative covariance between reproduction and survival. In the real world, however, the opposite patterns also can be found, and these patterns can be explained (and even expected) when one accepts the possibility that individuals are not interchangeable (van Noordwijk and de Jong 1986). Persistent individual differences in reproductive success (Lee et al. 2011) are generally taken to reflect individual differences in ‘quality’, which is a rather slippery concept but is generally thought to be positively correlated with fitness (Wilson and Nussey 2010). Individual differences in quality can be influenced by both genetic and environmental factors (Byholm et al. 2007) and can persist across many reproductive seasons [e.g., owing to long-lasting maternal effects (Mousseau and Fox 1998; Kruuk 2004) or mating dynamics (McElligot et al. 2000; Pelletier et al. 2006)]. In species for which fecundity increases with age (as applies to most ectotherms with indeterminate growth), persistent individual differences are likely the rule rather than the exception: individuals that are large for their age early in life will generally have relatively high reproductive success, and they also are likely to be large for their age in later years (Waples and Feutry 2022). Likewise, if individual quality is high for both reproduction and survival, that can offset any costs of reproduction and lead to a positive covariance (Smith 1981; Pelletier et al. 2006), a result that also can occur if mortality is anthropogenically modulated (e.g., if hunters avoid killing female moose with calves; Lee et al. 2020).

The second factor is that although monitoring evolutionary dynamics in age-structured populations remains logistically challenging, especially for long-lived species, the recent genomics revolution has greatly increased our ability to reconstruct population pedigrees using non-invasive genetic samples. Consider, then, the following scenario:

- A researcher is conducting a long-term pedigreed study of their focal species;
- The pedigree data are sufficient to robustly estimate generation length and variance in *LRS;*
- The biology of the focal species is such that it generates substantial autocorrelation of reproductive success and/or covariance of reproduction and survival.

A key question then becomes, “If the empirical estimates of *T* and *V*_*k*•_ are used in Equation 1 to estimate *N_e_*, will the result accurately reflect the rate of genetic drift in the population?”

The goal of this study is to answer this question. Using hypothetical vital rates for populations with overlapping generations, multi-year population pedigrees are simulated according to three Models that a) simulate independence of reproduction and survival over time (Model ‘NoCor’); b) generate strong positive autocorrelations and covariances (Model ‘Positive’)); and c) generate strong negative autocorrelations and covariances (Model ‘Negative’)). Each model is simulated under three levels of reproductive skew: Low, Medium, and High. Along each simulated pedigree, genetic variation is tracked at a number of loci, and the rates of genetic drift are quantified by two common metrics: rate of decline in heterozygosity, and rate of increase in allele frequency variance. These observed rates are compared with expected rates based on *N_e_* calculated from the population pedigree using Equation 1.

## METHODS

### Population demography

The core evaluations modeled reproduction and genetic change in a hypothetical population with age at maturity α=3 and maximum age ω=10, which produced a maximum adult lifespan of *AL*=8 years (ages 3-10, inclusive; see core vital rates in Table 1). Population dynamics followed the discrete time, birth-pulse model of Caswell (2001), where individuals that reach age *x* produce on average *b_x_* offspring and then survive to age *x*+1 with probability *s_x_*. Because previous work had evaluated sensitivity to the constant-size assumption, to limit the number of potentially confounding variables, age structure was fixed and defined by the vector of cumulative survival through age *x*, *l_x_*. Offspring were enumerated at age 1, so setting *l_1_*=1 and letting *N_1_* be the number of offspring in each cohort that reach age 1, the numbers in each successive age class (*x*=2,ω) are given by *N_x_*=*N_1_l_x._* For the core life table in Table 1, *N_1_*=200 and the full vector of age-class abundance was (rounded to the nearest integer): *N_x_*=[200,140,98,69,48,34,24,16,12,8]. Total abundance was Σ*N_x_*=649, of which 340 were juveniles, so the adult census size was *N_Adult_* = 309. These numbers apply to a single sex; sex ratio was 1:1 in the core analyses, so total population size was twice as large.

**Table 1.**
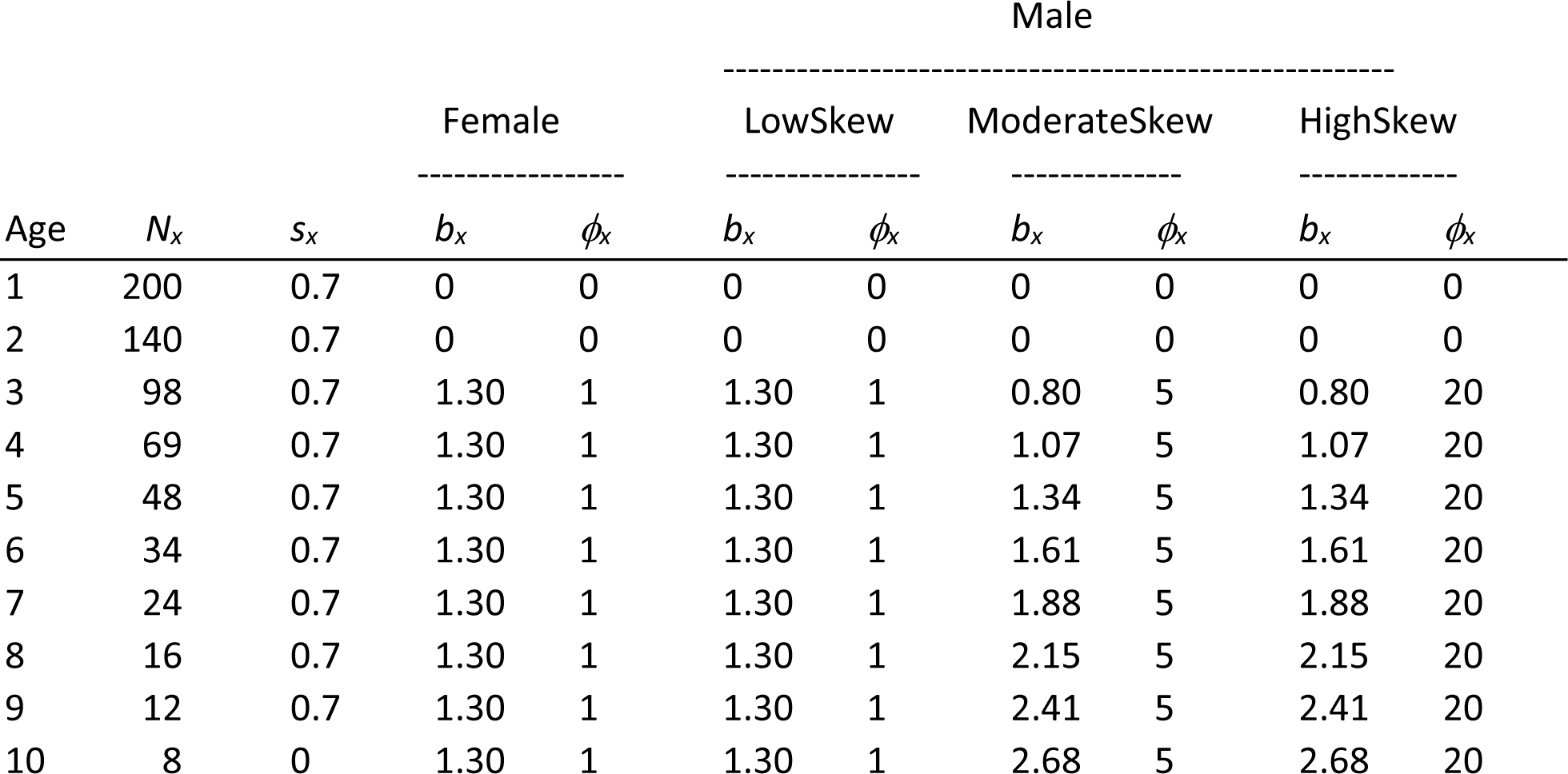
Core vital rates used in the simulations. The hypothetical population matures at age 3 and has a maximum age of 10. Survival is constant at *s_x_* = 0.7/year, and the vector *N_x_* is the number of individuals of each sex in each age class. In females, fecundity (*b_x_*) is always constant with age, and within each age the distribution of offspring number is Poisson (*ϕ*=1). In Scenario LowSkew, male vital rates are identical to those for females. In the other scenarios, male fecundity increases with age, and within ages reproductive variance is either substantially overdispersed (*ϕ*=5; Scenario ModerateSkew) or very strongly overdispersed (*ϕ*=20; Scenario HighSkew). Fecundity is scaled to values that will produce a stable population. Table S5 shows vital rates for some other life histories that were modeled.

The core life table allowed patterns of age-specific fecundity and reproductive skew to differ between the sexes. In all analyses, fecundity was constant with age in females and reproduction followed a Poisson process (all *ϕ*=1), so every year all females behaved like a single Wright-Fisher population with essentially random variation in offspring number. In males, three reproductive-skew levels were modeled. In LowSkew, *b_x_* and *ϕ_x_* were identical to values for females. In ModerateSkew, fecundity for males was proportional to age, and reproductive skew was moderately strong for all ages (*ϕ_x_*=5, indicating that variance in offspring number was 5 times the mean). In HighSkew, male *b_x_* was also proportional to age, and reproductive skew was very strong (all *ϕ_x_*=20). Fecundities were scaled to values that would produce a stable population by ensuring that Σ*b_x_l_x_*=2. With fecundities so scaled, the expected number of offspring produced in each time period by each age class within each sex was *B_x_*=*b_x_N_x_*, with Σ*B_x_*=2*N_1_*=400 total offspring, of which half are male and half female.

### Modeling reproduction

To ensure that the realized distribution of offspring number closely approximated the parametric values in the life table, an algorithm was developed (*NegBinom*) that used a negative-binomial simulator to generate random vectors of offspring numbers (***k***) expected to have the desired mean and variance. A disadvantage of this approach is that the realized mean of the simulated distribution is a random variable; as a consequence, the total number of offspring produced (Σ***k***) also varies randomly and only by chance equals the target number. To maintain a constant total number of offspring in each cohort, the following procedure was used:

- For each age *x*, the *rnbinom* function in R was used to generate *N_x_* random *k* values, using the parameterization *mu*=*b_x_* and *size*=*b_x_*^2^/(*V_k(x)_*-*b_x_*), with *V_k(x)_*=*ϕ_x_b_x_*. This function requires *V_k(x)_*>*b_x_*, so for LowSkew scenarios with *V_k(x)_*=*b_x_* the *rpois* function was used instead.
- The total length of all the age-specific ***k*** vectors was compared with the target cohort size = 2*N_1_*. This process was repeated until the total length fell in the range [2*N_1_*, 1.05*2*N_1_*]. At that point, the vectors of offspring numbers were converted to a list of parental IDs and this list was randomly downsampled (if necessary) to reach exactly 2*N_1_* offspring.
- This entire process was repeated for the opposite sex. Two related issues require consideration:
- Does reproduction in one time period affect reproduction in any subsequent time period?
- Does reproduction in one time period affect the probability of survival to subsequent time periods?

To see the consequences of a positive answer to either of these questions, note that the expected lifetime reproductive success [*E*(*k•*)] of an individual is a simple additive function of its age-specific fecundity and its age at death (*q*):

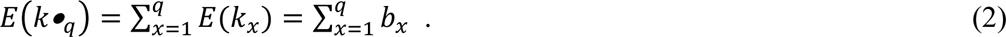

The variance of a sum, however, also depends on the covariance structure of the terms being added: var(*A*+*B*)=var(*A*)+var(*B*)+2cov(*A*,*B*). More generally, when applied to the variance of a sum of *q* terms:

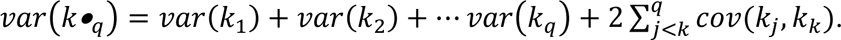

Since *var*(*k*_*x*_) = *ϕxb*_*x*_, the above can be written as

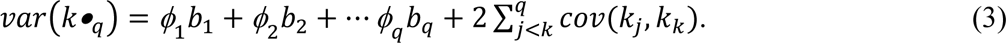

Equation 3 gives the variance in offspring number among individuals that die at a single age (*q*). The lifetime variance in offspring number among all individuals in a cohort can be obtained using the definition of a variance as *E*(*x*^2^) – [*E*(*x*)]^2^. In the current notation,

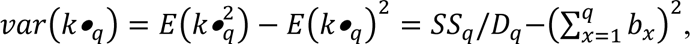

where *D*_*q*_ is the number of individuals that die after reaching age *q* and 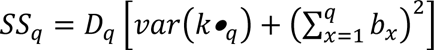 is the sum of the squared offspring numbers for these *D*_*q*_ individuals. If we ignore individuals that die before reaching age at maturity, *Σ*(*D*_*q*_) is the number of individuals in each cohort that reach adulthood. The total sums of squares is obtained by summation: *SS*_*T*_ = Σ(*SS*_*q*_), and the lifetime variance in offspring number is calculated as

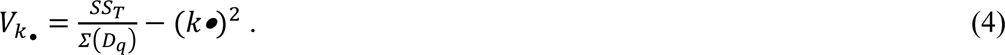

The *AGENE* model (Waples et al. 2011) calculates *V*_*k*•_ and *N_e_* using age-specific vital rates from an expanded life table and assuming that expected values of all the covariance terms are zero.

Developing analytical expectations for the covariance terms when they cannot be assumed to be zero is not a simple task, especially in any generalized form. However, the simulation algorithm can be tweaked to generate positive or negative correlations in reproductive success, as can be illustrated with a hypothetical example involving lifetime reproductive success of a cohort of 20 individuals that is tracked for a 5-year maximum lifespan (Table 2). *NA*s in the table indicate individuals that were not alive at the specified age. In this example, 8 individuals died after age 1 and before reaching age 2, 4 died after age 2, and 3 each died after ages 3 and 4, leaving just 2 individuals that survived to age 5. Expected fecundity increased linearly with age (*b_x_* = [1,2,3,4,5]), and the parametric within-age variance was 5 times the mean (ModerateSkew; *ϕ*=5) for each age. To populate the table, vectors of offspring numbers were randomly generated for each age using the *rnbinom* function in R (R core team 2022) (generating 20 random values for age 1 with mean = 1 and variance = 5; 12 random values for age 2 with mean = 2 and variance = 10; etc.).

**Table 2.**
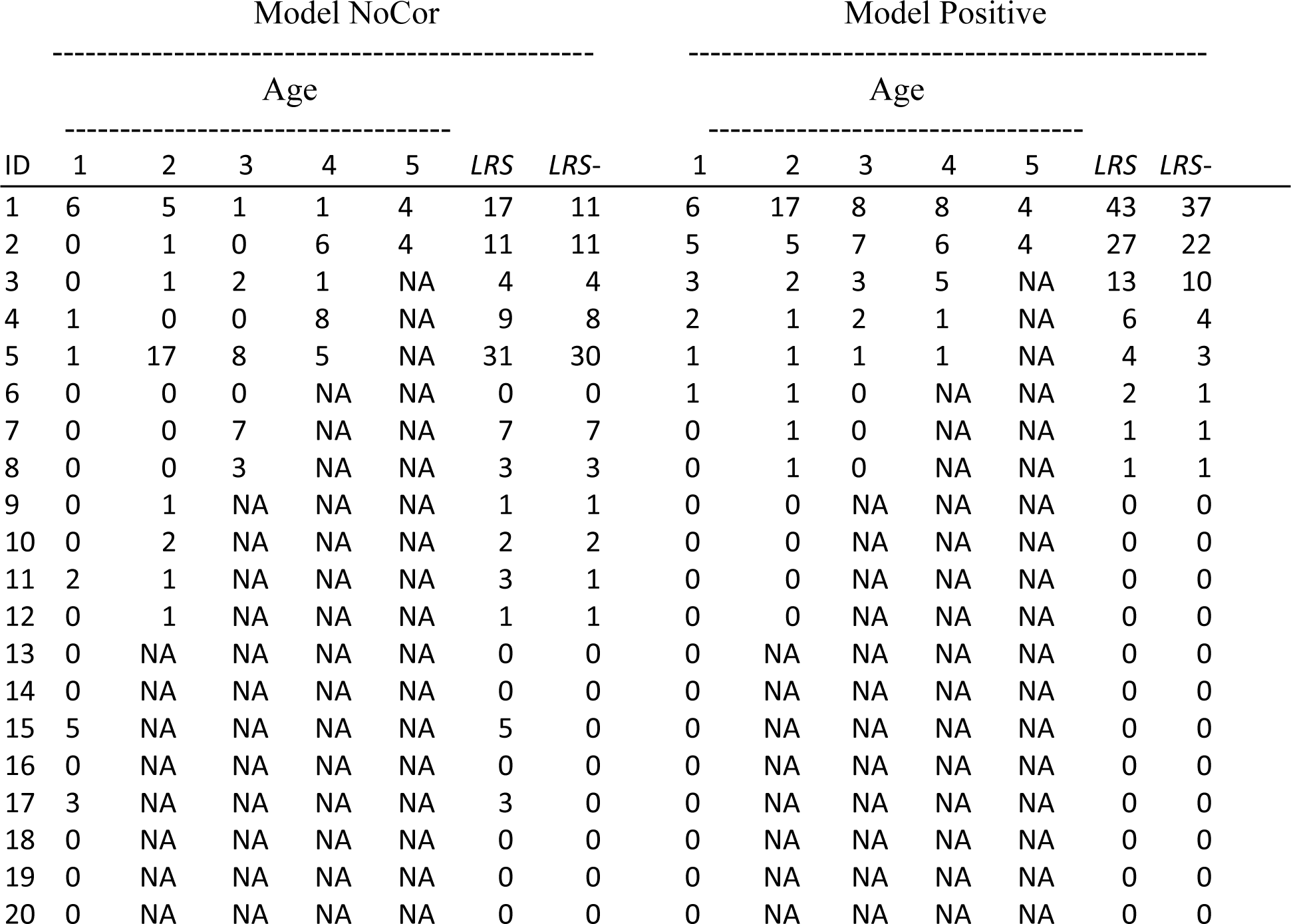
Example illustrating how positive correlations in reproductive success over time were generated in simulated populations. For each of 20 individuals in a cohort, the number of offspring produced at each age 1-5 is shown in the left table. Each year individuals with the highest ID numbers die, after which time their reproductive success is recorded as NA. In the left set of columns, offspring numbers were randomly generated for survivors at each age, as occurs under Model NoCor; in the right set of columns, those random numbers were sorted so that each year the individual with the lowest ID produced the most offspring (as occurs in Model Positive). The columns labeled ‘*LRS*’ show the total lifetime numbers of offspring produced by each individual, and the columns labeled ‘*LRS*-’ show *LRS* for all ages except age 1. See text for discussion and Table S1 for an example illustrating Model Negative.

| ID | Model NoCor |  |  |  |  |  |  | Model Positive |  |  |  |  |  |  |
| --- | --- | --- | --- | --- | --- | --- | --- | --- | --- | --- | --- | --- | --- | --- |
|  | ----- |  |  |  |  |  |  | ----- |  |  |  |  |  |  |
|  | Age |  |  |  |  |  |  | Age |  |  |  |  |  |  |
|  | ----- |  |  |  |  |  |  | ----- |  |  |  |  |  |  |
|  | 1 | 2 | 3 | 4 | 5 | *LRS* | *LRS-* | 1 | 2 | 3 | 4 | 5 | *LRS* | *LRS-* |
| 1 | 6 | 5 | 1 | 1 | 4 | 17 | 11 | 6 | 17 | 8 | 8 | 4 | 43 | 37 |
| 2 | 0 | 1 | 0 | 6 | 4 | 11 | 11 | 5 | 5 | 7 | 6 | 4 | 27 | 22 |
| 3 | 0 | 1 | 2 | 1 | NA | 4 | 4 | 3 | 2 | 3 | 5 | NA | 13 | 10 |
| 4 | 1 | 0 | 0 | 8 | NA | 9 | 8 | 2 | 1 | 2 | 1 | NA | 6 | 4 |
| 5 | 1 | 17 | 8 | 5 | NA | 31 | 30 | 1 | 1 | 1 | 1 | NA | 4 | 3 |
| 6 | 0 | 0 | 0 | NA | NA | 0 | 0 | 1 | 1 | 0 | NA | NA | 2 | 1 |
| 7 | 0 | 0 | 7 | NA | NA | 7 | 7 | 0 | 1 | 0 | NA | NA | 1 | 1 |
| 8 | 0 | 0 | 3 | NA | NA | 3 | 3 | 0 | 1 | 0 | NA | NA | 1 | 1 |
| 9 | 0 | 1 | NA | NA | NA | 1 | 1 | 0 | 0 | NA | NA | NA | 0 | 0 |
| 10 | 0 | 2 | NA | NA | NA | 2 | 2 | 0 | 0 | NA | NA | NA | 0 | 0 |
| 11 | 2 | 1 | NA | NA | NA | 3 | 1 | 0 | 0 | NA | NA | NA | 0 | 0 |
| 12 | 0 | 1 | NA | NA | NA | 1 | 1 | 0 | 0 | NA | NA | NA | 0 | 0 |
| 13 | 0 | NA | NA | NA | NA | 0 | 0 | 0 | NA | NA | NA | NA | 0 | 0 |
| 14 | 0 | NA | NA | NA | NA | 0 | 0 | 0 | NA | NA | NA | NA | 0 | 0 |
| 15 | 5 | NA | NA | NA | NA | 5 | 0 | 0 | NA | NA | NA | NA | 0 | 0 |
| 16 | 0 | NA | NA | NA | NA | 0 | 0 | 0 | NA | NA | NA | NA | 0 | 0 |
| 17 | 3 | NA | NA | NA | NA | 3 | 0 | 0 | NA | NA | NA | NA | 0 | 0 |
| 18 | 0 | NA | NA | NA | NA | 0 | 0 | 0 | NA | NA | NA | NA | 0 | 0 |
| 19 | 0 | NA | NA | NA | NA | 0 | 0 | 0 | NA | NA | NA | NA | 0 | 0 |
| 20 | 0 | NA | NA | NA | NA | 0 | 0 | 0 | NA | NA | NA | NA | 0 | 0 |

Results of one random realization of this process are shown in the left side of Table 2. At age 1, 2 individuals produced exactly 1 offspring, single individuals produced 2,3,5, and 6 offspring each, and the remaining 14 individuals produced no offspring. The ‘*LRS*’ column shows that across the original cohort, lifetime offspring number ranged from 0 (6 individuals) to 31 (individual 5). Three factors contribute to the variance in *LRS* in this example. 1) Individuals that (by luck or pluck) survive to older ages have more opportunities to add to their *LRS*. 2) Because fecundity increases with age in this example, longer-lived individuals get an additional bonus because their reproductive success is higher in later years. 3) Variance in reproductive success is overdispersed within each age (*ϕ*>1). Collectively, these factors cause lifetime *V*_*k*•_ (58.1) to be much higher than the mean (4.85). So far, this example has not implemented any covariance of survival and reproduction or any temporal autocorrelations in individual reproductive success over time. We refer to this as Model NoCor.

Temporal correlations in reproductive success are easy to generate by sorting the randomly-generated vectors of offspring number before mapping them to individuals. The right side of Table 2 shows the results of sorting the ***k*** vector for each age such that the largest value is assigned to individual 1, the next largest to individual 2, and so on. This simple ploy accomplishes two things: 1) It creates persistent individual differences in reproductive success, which manifest as positive correlations between an individual’s reproductive output across time; 2) It creates positive correlations between reproduction and subsequent survival, which enhance the strength of the persistent individual differences. This is Model Positive. The net result is that *V*_*k*•_ more than doubles (to 122.9) while the mean remains the same.

In empirical datasets, the pairwise covariance terms can be challenging to deal with because a) they are very numerous for long-lived species, and b) sample sizes are generally small for comparisons involving older age classes. Here a new metric is introduced (ρ*_α_*,*_α_***_+_**) that generally can be applied to all individuals in a cohort that reach age at maturity. This metric represents the Pearson correlation coefficient between two vectors: ***k_α_*** = offspring number for all individuals at the age at first reproduction (α), and ***k*_α+_** = *LRS* - ***k_α_*** = lifetime reproductive success of the same individuals for all subsequent years. In the example for Model NoCor in Table 2, this correlation is slightly positive (ρ*_α_*,*_α_***_+_**=0.14) but not significantly so (*P*>0.5 for a two-tailed test). For the extreme Model Positive, in contrast, this correlation was close to unity (ρ*_α_*,*_α_***_+_**=0.96; *P*<0.001).

Negative temporal correlations in reproductive success are easy to generate by reversing the sorting process and assigning the largest *k* value each year to the individual with the highest ID number (Model Negative; see Figure S1). Since individuals with the highest ID numbers are the ones that die each year after reproduction, this ensures that new individuals get to reproduce each year, which in turn reduces *V*_*k*•_. Applying Model Negative to the simulated data in Table 2 reduced *V*_*k*•_ sharply (to 17.4) and led to a significantly negative correlation (ρ*_α_*,*_α_***_+_**=-0.43; *P*<0.05; Table S1).

To illustrate an alternative way to generate correlations between reproduction and survival, the analyses in the main text were repeated using a second simulation algorithm. *TheWeight* (Waples 2020, 2022a) is a generalized Wright-Fisher model that allows for unequal parental expectations of reproductive success, specified by a vector of parental weights, ***W***. Details for how this algorithm was implemented are in Supporting Information.

It is worth noting that Equation 3 for *var*(*k•*_*q*_) (and by extension Equation 4 for *V*_*k*•_) do not contain any terms for the covariance of individual reproduction and survival. To the extent these covariances are non-zero, they can provide insights into key evolutionary processes. With respect to variance in reproductive success, however, any effects of these covariances manifest themselves as positive or negative autocorrelations in individual reproductive success over time, so for the analysis of *V*_*k*•_ it is sufficient to focus on these autocorrelation terms.

### Tracking genetic drift

Table 2 illustrates how lifetime *V*_*k*•_ was calculated from simulated demographic data. To determine whether Hill’s *N_e_* based on these *V*_*k*•_ values accurately predicted the rate of genetic drift, two common genetic metrics were monitored. The expected variance in allele frequency (*V*_*p*(*t*)_) after *t* generations of genetic drift is (Hedrick 2000):

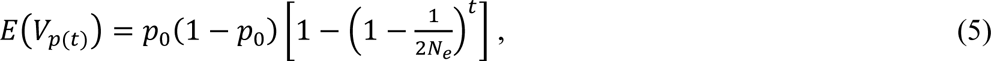

where *p*_0_ is the initial allele frequency. In isolated populations with no mutation, random changes in allele frequency also cause an increase in homozygosity over time, such that after *t* generations the expected amount of remaining heterozygosity (*H*_*t*_) is a simple function of initial average heterozygosity (*H*_0_) and *N_e_* (Crow and Kimura 1970):

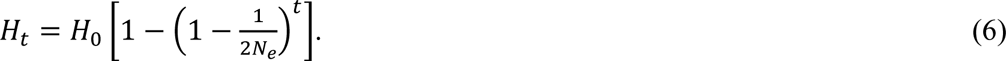

To monitor these metrics, genetic variation was tracked at *L* unlinked, diallelic (∼SNP) loci. Genotypes were recorded as [0,1,2], indicating the number of copies of the focal allele each individual carried. In each replicate simulation, the population was initialized by filling each age class with the appropriate number (*N_x_*) of individuals. In year 0, all individuals were designated as heterozygotes (genotype ‘1’) at every locus, so initial allele frequencies were all *p*_0_ = 0.5. In year 1 and subsequent years, mean observed *H*_*t*_ and *V*_*p*(*t*)_ were computed for all members of the newborn cohort. As a single episode of random mating is sufficient to establish Hardy-Weinberg genotypic ratios, mean *H* at year 1 was on average 0.5. Years were converted into generations using the relationship *t*=*y*/*T*, where *y* is elapsed time in years (*T* = generation length was 4.84 for Scenario LowSkew and 5.22 for the other scenarios where male fecundity increased with age.)

Observed rates of genetic drift were compared with expected rates calculated two ways. Under Model NoCor, where probabilities of reproduction and survival are independent over time, the expected variance in *LRS* and hence Hill’s *N_e_* can be calculated from an expanded life table (age-specific survival, fecundity, and *ϕ*) using the AgeNe model (Waples et al. 2011). The resulting *N_e_* was then used in Equations 5 and 6 to generate expected values for the two genetic drift indices. The second approach used the population pedigree from the simulations to calculate *V*_*k*•_ for each annual cohort of offspring, and from this a ‘pedigree’ *N_e_* was calculated every year using Equation 1. The harmonic mean pedigree *N_e_* was then used in Equations 5 and 6 to predict expected rates of genetic drift based on the actual population pedigree.

With age structure, the rate of increase in *V*_*p*(*t*)_ reaches a steady state only after a burnin period lasting several generations, which means that Equation 5 might not be accurate in the early years. To account for this effect, after the burnin period, the empirical *V*_*p*(*t*)_ for year 50 was averaged across replicates to produce *V*_*p*(*Burnin*)_, and the subsequent increase in *V*_*p*_ with time was calculated as *ΔV*_*p*(*t*)_ = *V*_*p*(*t*)_ − *V*_*p*(*Burnin*)_. With this adjustment, *E*(*ΔV*_*p*(*t*)_) can be calculated from Equation 5, replacing *p*_0_(1 − *p*_0_) with the mean *p*_50_(1 − *p*_50_) averaged across loci and replicates at the end of the burnin period.

Each replicate simulation was run for 500 years. Except as noted, results shown here are averaged across 10 replicate simulations, each tracking genetic variation at *L*=100 loci.

## RESULTS

The two simulation algorithms were both successful in achieving the desired level of reproductive skew and covariances/autocorrelations. For simplicity, results for *NegBinom* are presented in the main text and those for *TheWeight* are in Supporting Information.

### Model NoCor

#### Population demography

For Model NoCor, analytical expectations for lifetime *V*_*k*•_ and *N_e_* are possible based on the vital rates in Table 1, and these provide a useful reference point for evaluating results of the simulations. Scenario LowSkew is the simplest as fecundity is constant in both sexes, with *ϕ*=1 for all ages. This means that all adults reproducing each year behave like a single Wright-Fisher population with Poisson variation in reproductive success. Under this scenario, the parametric expectation for *V*_*k*•_ is 10.2 (Table 3). Poisson variance in *LRS* would lead to *V*_*k*•_ equal to the mean (which must be 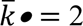 in a stable population), so most of the total lifetime variance can be attributed to variation in longevity (some individuals live longer than others and have more opportunities to reproduce). In Scenario ModerateSkew, male fecundity increases with age, and at each age the variance in male offspring number is 5 times the mean (*ϕ*=5). Together these factors increase parametric *V*_*k*•_ to 16.1 (Table 3). In Scenario HighSkew, *ϕ* takes a rather extreme value of 20, and parametric *V*_*k*•_ grows to 31.1 – over 15 times the lifetime mean.

**Table 3.** Summary of simulation results for three reproductive-skew Scenarios (LowSkew→ϕ=1; ModerateSkew→ϕ=5; HighSkew→ϕ=20). For each Scenario, the parametric *V*_*k*•_ and *N_e_* under Model NoCor are shown. Columns on the left show the *N_e_* calculated from the simulated pedigrees; middle columns show the ratio of observed heterozygosity at generation *t* to the expected value based from Equation 6 using the pedigree *N_e_*; columns on the right show the correlation (ρ*_α_*,*_α_***_+_**) between offspring numbers for each individual at age α and for all subsequent ages combined (as shown in the *LRS-* column from Table 2). Pedigree *N_e_* was calculated as the harmonic mean across cohorts and replicates; the ratio Obs/Exp *H_t_* was the average across 10 replicates, and within each replicate the ratio was calculated as the mean across the final 50 years. For each cohort in each replicate, ρ*_α_*,*_α_***_+_** was computed across all individuals that reached age at maturity, and results were averaged across cohorts and replicates. See Table S4 for comparison of observed and expected rates on increase in allele frequency variance, *V*_*p*(*t*)_.

| Scenario | Model NoCor | | Pedigree $N_e$ | | | Obs/Exp $H_t$ | | | Correlation ($\rho_{a,\alpha+}$) | | | |
| --- | --- | --- | --- | --- | --- | --- | --- | --- | --- | --- | --- | --- |
|  | Model |  | Model |  |  | Model |  |  | Model Positive |  | Model Negative |  |
| | $E(V_{k\bullet})$ | $E(N_e)$ | NoCor | Positive | Negative | NoCor | Positive | Negative | F | M | F | M |
| LowSkew | 10.2 | 636 | 640 | 324 | 999 | 0.999 | 1.007 | 1.003 | 0.85 | 0.85 | -0.68 | -0.69 |
| ModerateSkew | 16.1 | 462 | 468 | 159 | 795 | 0.999 | 1.008 | 0.999 |  | 0.94 |  | -0.34 |
| HighSkew | 31.1 | 253 | 270 | 72 | 367 | 0.996 | 1.015 | 0.999 |  | 0.94 |  | -0.11 |

These demographic changes have predictable consequences for effective population size under Model NoCor. In Scenario LowSkew, where both sexes have identical vital rates (leading to *T*=4.84 and *V*_*k*•_ = 10.2), parametric *N_e_* from Equation 1 is 4*400*4.84/(2+10.18) = 636. In the scenarios with moderate to high skew, increasing fecundity with age increased generation length in males from 4.84 to 5.59, so across both sexes overall *T* was 5.21. All else being equal, *N_e_* increases linearly with generation length (Equation 1). However, oversdispersion in male reproductive success substantially increased in these scenarios, and this more than offset the modest increase in generation length. The parametric expectations for *N_e_* are 452 for Scenario ModerateSkew and 253 for Scenario HighSkew (Table 3), which are, respectively, 29% and 60% lower than for Scenario LowSkew.

The new correlation metric, ρ*_α_*,*_α_***_+_**, examines the association between an individual’s reproductive success in the first year of sexual maturity (age 3 for the core life table) and the rest of its life. As expected under Model NoCor, mean values of ρ*_α_*,*_α_***_+_** for both sexes for all three scenarios were close to zero, all falling in the range [−0.01, +0.01] (data not shown).

Mean empirical *V*_*k*•_ in the NoCor simulations agreed well with the parametric expectations (Figure 1). Harmonic mean *N_e_* calculated from the simulated pedigrees was within 1% of the parametric expectation for Scenarios LowSkew and ModerateSkew and ∼7% higher for Scenario HighSkew (Table 3). This latter result reflects the difficulty in precisely modeling strongly overdispersed variance in reproductive success, especially when older age classes have few individuals (only 8 of each sex for the oldest age class in the core life table).

**Figure 1.**
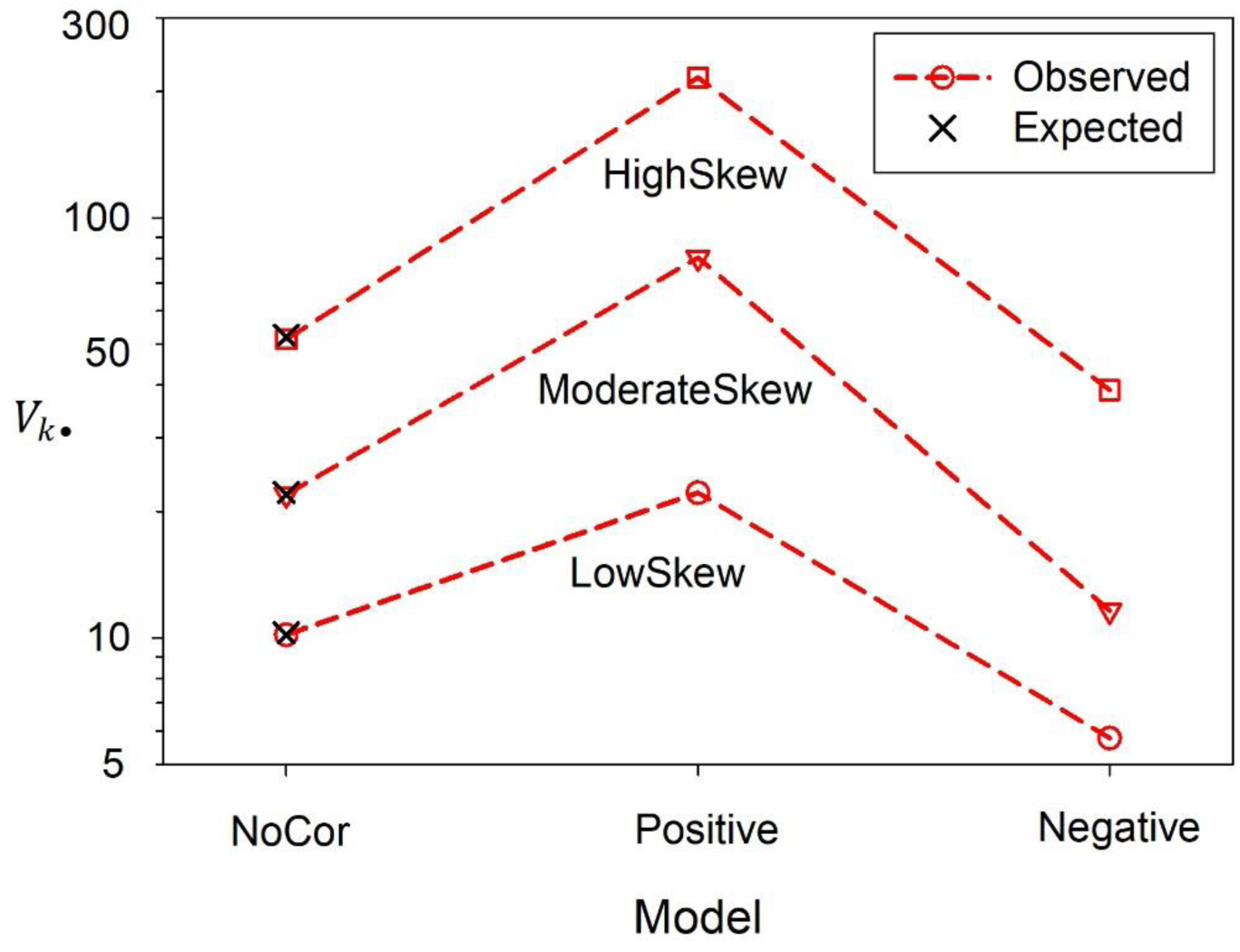
Mean values of the lifetime variance in reproductive success (*V*_*k*•_) for simulated data. Results are shown for the three Models and three reproductive-skew Scenarios described in the text. Parametric expectations based on the vital rates in Table 1, which are only available for Model NoCor, are shown by black Xs. Note the log scale on the *Y* axis.

#### Tracking genetic drift

For all 3 Scenarios under Model NoCor, the mean ratio of observed to expected heterozygosity calculated over the last 50 years of each replicate was close to unity (all values within 1% of 1.0; Table 3). A comparable result was found for comparisons of observed and expected variance in allele frequency (Table S4). These results are consistent with but extend previously reported results for Hill’s model. Waples et al. (2014) found excellent agreement between observed and expected rates of decline in heterozygosity in simulations based on vital rates for 20 different species, but for most species it was assumed that *ϕ*=1 for all ages. Results for Scenario HighSkew, with very high within-age reproductive skew (*ϕ*=20), are therefore new.

### Models with Correlations

#### Population demography

The realized annual offspring numbers each year were sorted in Models Positive and Negative. Although these offspring numbers spanned a relatively small range of values for Scenario LowSkew, when sorted before assigning to individuals they had a substantial effect on population demography. For Scenario LowSkew, *V*_*k*•_ more than doubled under Model Positive and was nearly halved under Model Negative (Figure 1), with corresponding changes to pedigree *N_e_* (Table 3), and ρ*_α_*,*_α_***_+_** was strongly positive in both sexes (0.85) for Model Positive and strongly negative in both sexes (−0.68 to −0.69) for Model Negative (Table 3).

With stronger reproductive skew, results were even more dramatic. For Model Positive, ρ*_α_*,*_α_***_+_** was >0.9 (Table 3). These strong positive correlations concentrated reproduction in just a few individuals, which in turn substantially increased lifetime variance in reproductive success. With *ϕ*=5 (Scenario ModerateSkew), lifetime *V*_*k*•_ increased almost four-fold, and with *ϕ*=20 (Scenario HighSkew), *V*_*k*•_ more than quadrupled, to >200 (Figure 1). Increases in *V*_*k*•_ caused corresponding decreases in effective size (Table 3). For Scenario ModerateSkew, *N_e_* was less than half of the parametric value expected under Model NoCor, and for Scenario HighSkew realized *N_e_* was less than 30% of the value expected Model NoCor.

In Model Negative, individuals who were assigned the largest numbers of offspring each year all died before reaching the next age. This created a strong negative correlation between initial and subsequent reproductive success: ρ*_α_*,*_α_***_+_** = −0.34 for Scenario ModerateSkew and −0.11 for Scenario HighSkew (Table 3). These negative correlations minimized disparities in lifetime reproductive success and reduced *V*_*k*•_ compared to expectations under the NoCor model (Figure 1) and consequently increased effective size (Table 3).

#### Tracking genetic drift

Even for extreme versions of correlated reproduction, use of the pedigree *N_e_* in Hill’s Equation 1 accurately predicted the rates of loss of heterozygosity and increase in allele frequency variance (Tables 3 and S2; Figures 2 and 3). For loss of heterozygosity, all deviations from expectations were <1% except for the extremely overdispersed (*ϕ* =20) Scenario HighSkew under Model Positive, where mean heterozygosity in the last 50 years was 1.5% higher than expected (Table 3). Stochastic variation in the rate of increase in allele frequency variance was somewhat higher, but for all Scenario x Model combinations the observed change in *V*_*p*(*t*)_ was within a few % of the expected (Table S4), with an overall mean observed/expected ratio of 1.007.

**Figure 2.**
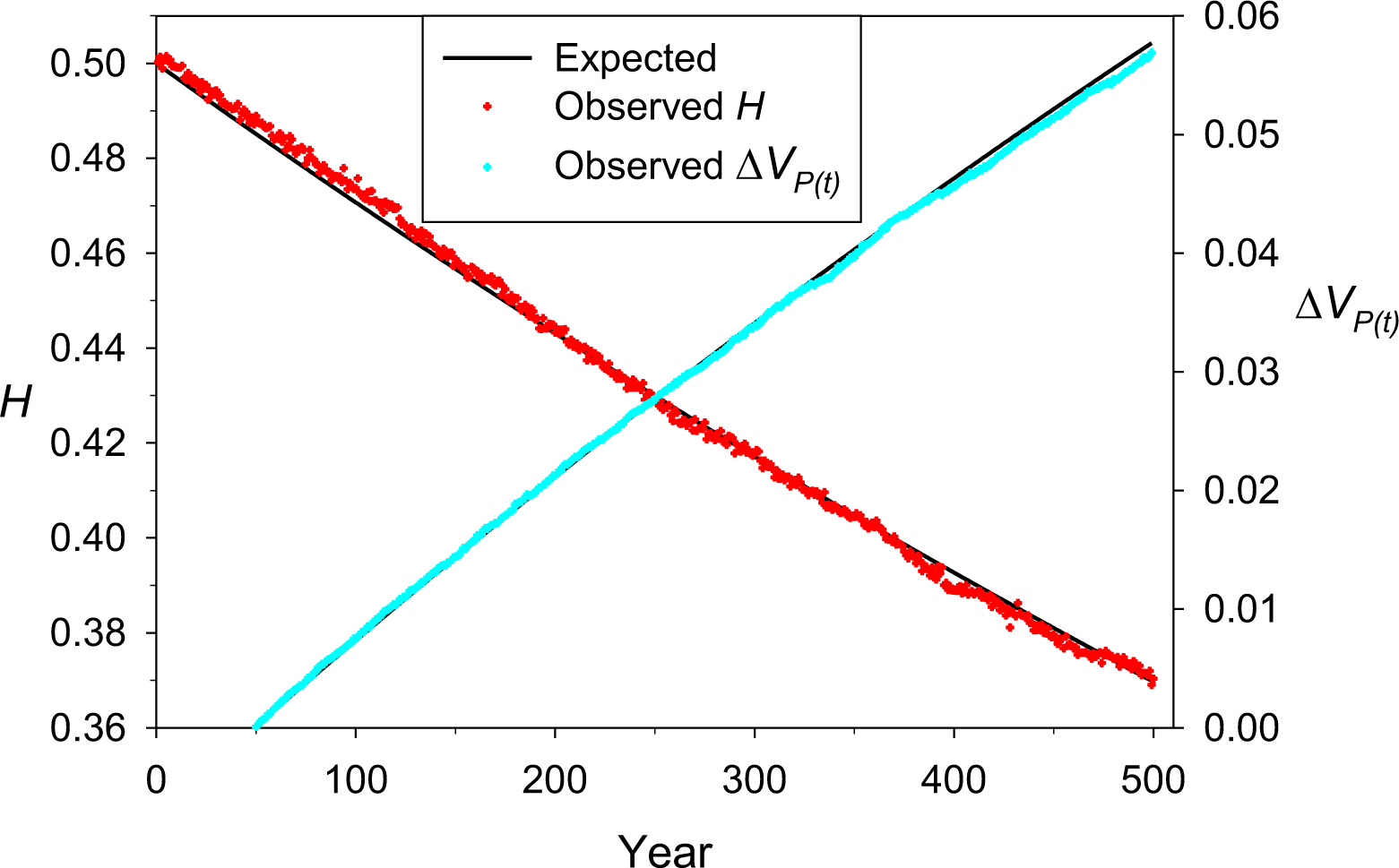
Observed (colored symbols) and expected (black lines) rates of genetic drift in a simulated population under Model Positive, Scenario ModerateSkew (in which *ϕ*=5 for all ages in males). The left *Y* axis and red symbols plot loss of heterozygosity and the right *Y* axis and cyan symbols plot increasing variance in allele frequency. Expected rates of genetic drift were computed using Equations 5 and 6 and the harmonic mean pedigree *N_e_* for this scenario (which was 158). Results shown are averaged across 10 replicate 500-year pedigrees, each of which tracked genetic variation at 100 unlinked, diallelic loci.

**Figure 3.**
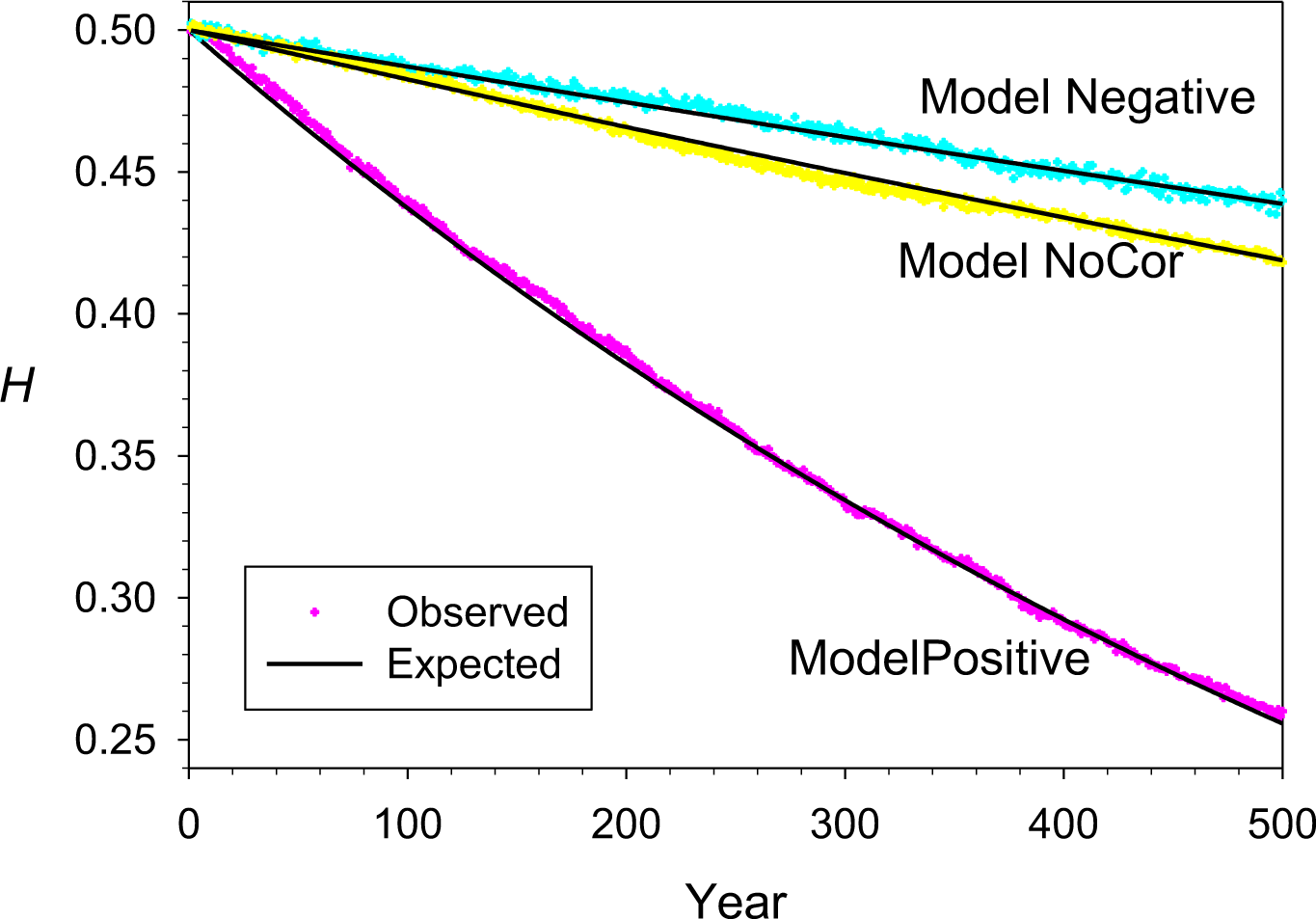
Observed (colored symbols) and expected (black lines) rates of loss of heterozygosity in simulated populations for three Models that lead to positive correlations (Model Positive), negative correlations (Model Negative), and independence of individual reproductive success over time (Model NoCor). These results are for Scenario HighSkew, in which ϕ = 20 for males of all ages. Results shown are averaged across 10 replicate 500-year pedigrees, each of which tracked genetic variation at 100 unlinked, diallelic loci.

### Alternate life histories

Results so far have all used variations of the vital rates in Table 1, which apply to a hypothetical species with 10 age classes. Simulations were also conducted for shorter lifespans (5 years, with age at maturity 1) and longer lifespans (20 years, with age at maturity 5) (Table S5). In both cases, fecundity was constant with *ϕ*=1 in females, and fecundity was proportional to age with *ϕ*=5 in males. As shown in Figure S1, Equation 1 based on *N_e_* calculated from the actual pedigrees accurately predicted the rates of loss of heterozygosity and increase in allele frequency variance for these different life histories.

### Precision

As the main focus of this paper is to evaluate potential bias in Equation 1 when it is applied to extreme demographic scenarios, a great deal of replication has been used to smooth out random demographic and genetic stochasticity to produce mean results that are qualitatively repeatable. Empirical datasets, on the other hand, generally are collected from a single realized population pedigree and might include data for a relatively small number of genes. As a reminder to researchers evaluating such empirical datasets, an example is included that generated a single 500-year population pedigree and tracked decline of heterozygosity in 10 different sets of 50 unlinked diallelic loci (Figure 4). At year 500 the average heterozygosity across the total 500 loci was close to the expected value from Equation 6 using the realized pedigree (0.256), but in one set of 50 loci mean (*H_obs_*) was >0.3 and in another set it was <0.2. Comparable results for allele frequency variance are shown in Figure 1 of Waples (2022b).

**Figure 4.**
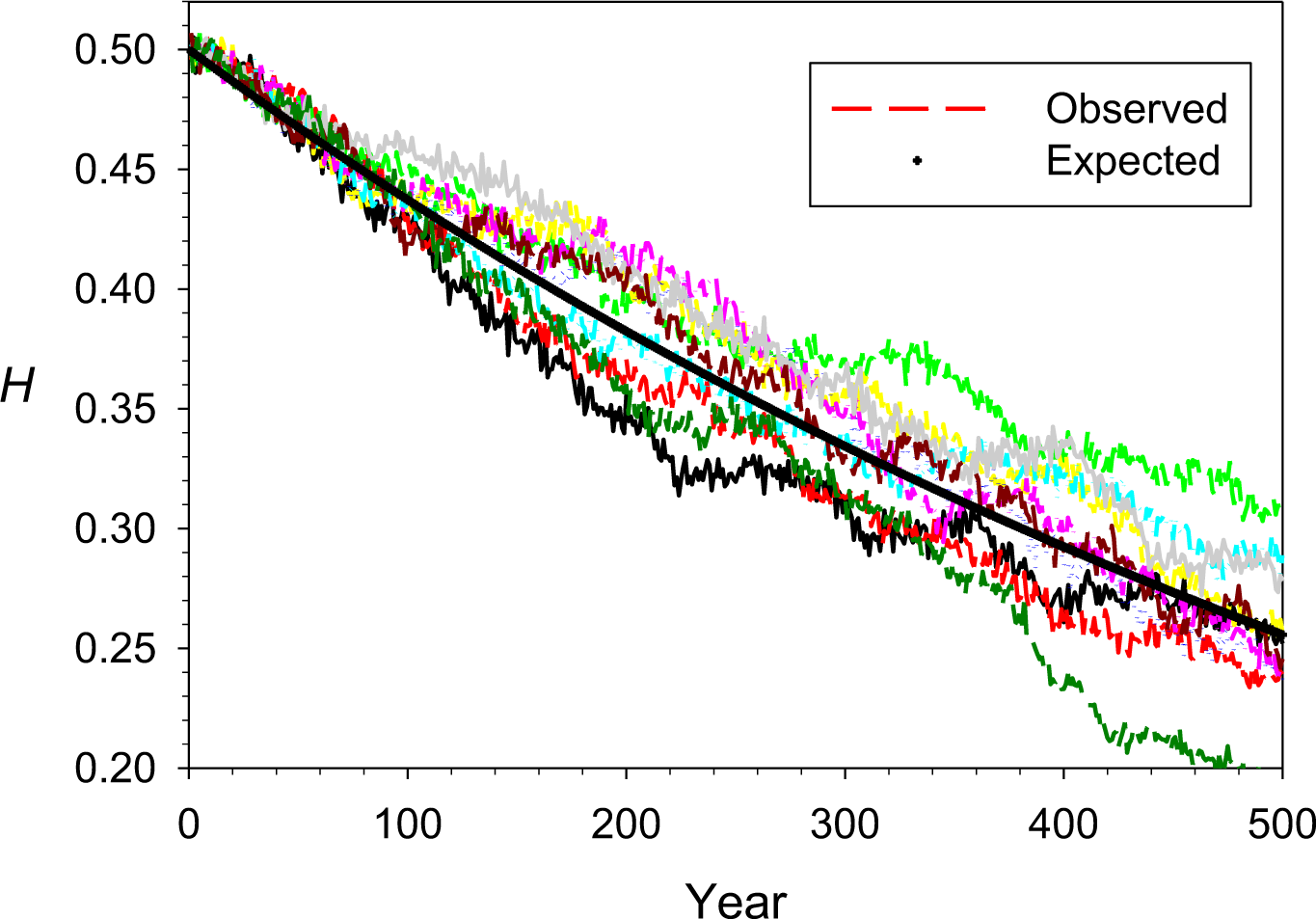
Observed (colored lines) and expected (solid black line) decline in observed heterozygosity for 10 different sets of 50 diallelic loci tracked on a single, 500-year pedigree. These results are for Model Positive and Scenario HighSkew, in which ϕ = 20 for males of all ages.

## DISCUSSION

Hill’s (1972) method for calculating effective population size is surprisingly robust to extreme reproduction scenarios. As expected, introducing strong autocorrelations in reproduction and covariance between reproduction and survival caused dramatic changes in lifetime variance in reproductive success. For Scenario LowSkew, mean *V*_*k*•_ was 3.8 times as large under Model Positive (positive autocorrelations and covariances) as it was under Model Negative (negative autocorrelations and covariances). For the scenarios that included substantial overdispersion of within-age reproductive success, the proportional differences were even greater (7.7 and 6.4 times larger for Model Positive for Scenarios ModerateSkew and HighSkew, respectively). The autocorrelations and covariances that arose when implementing Models Positive and Negative did not affect generation length, so when the empirically-derived estimates of *V*_*k*•_ were inserted in Hill’s Equation 1, they also led to realized effective sizes that differed dramatically among the three Models (Table 3).

The most important result from this study is that, when *V*_*k*•_ is computed from the population pedigree, *N_e_* calculated from Equation 1 accurately predicts the realized rate of genetic drift when inserted in Equations 5 (for rate of increase in allele frequency variance) and 6 (for rate of loss of heterozygosity). Excellent agreement between observed and predicted rates of genetic drift was found for diverse life histories (5, 10, and 20 year lifespans, with age at maturity 1, 3, or 5 years), for identical or different vital rates for males and females, and for extreme skew in reproductive success (variance up to 20 times the mean), all across nearly 100 generations of evolution.

These results are good news for researchers. Random processes in age-structured populations create dynamic heterogeneity in survival and reproduction (Vindenes et al. 2008; Tuljapurkar et al. 2009), and these processes are implemented here as Model NoCor. But the biological attributes of many species create autocorrelations of reproduction and/or covariances in reproduction and survival. As implemented here, Models Positive and Negative are more extreme than are likely to be found in most real species, but that was intentional. In the worst-case scenarios found here, observed rates of genetic drift were still within a few percent of those expected, and these scenarios involved very strong reproductive skew within ages. For all realistic applications to natural populations, therefore, Equation 1 can be considered to be a very reliable predictor of effective population size.

Although pairwise correlations of individual reproductive success in different years can provide valuable and detailed information regarding reproductive tradeoffs, the new index introduced here (ρ*_α_*,*_α_***_+_**) provides what appears to be a robust summary across the full lifespan. ρ*_α_*,*_α_***_+_** is the correlation between two vectors, one listing the number of offspring produced by each individual at the first age of sexual maturity (α) and the other listing the total number of offspring produced by the same individuals during the rest of their lifetimes. As expected, these correlations averaged close to 0 under Model NoCor and were consistently very high under Model Positive. Under Model Negative these average correlations were consistently negative, with a magnitude that depended on the degree of within-age reproductive skew. An advantage of this summary index compared to pairwise correlations of reproduction at specific ages is that the length of the two vectors considered by ρ*_α_*,*_α_***_+_** is the number of individuals in the cohort, rather than the inevitably smaller (and variable) number that survive to later ages. This increases statistical power, so ρ*_α_*,*_α_***_+_** might be used initially for diagnostic purposes before focusing in more detail on specific ages.

Steinar Engen and colleagues (Engen et al. 2005, 2009) have developed an alternative way to calculate *N_e_* when generations overlap and Hill’s (1972, 1979) assumptions of constant *N* and stable age structure are not met. In their model, the overall variance in *N* or total reproductive value over time arises from two additive components: an environmental variance 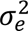, which quantifies effects of fluctuating environments over time, and a demographic variance 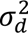, which quantifies effects of random demographic stochasticity within one time period, during which the environment is constant. Engen’s model uses a Leslie matrix to relate 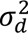 to a population’s age-specific vital rates (probabilities of reproduction and survival), and this allows formal consideration of the covariance of an individual’s fecundity at time *t* and that individual’s survival to time *t*+1. However, the time horizon for considering demographic stochasticity is only one time period, and Engen’s model does not include a term for lifetime variance in offspring number, so direct comparison with Hill’s model is only possible for some very simplified scenarios. The focus on a single time step means that Engen’s model cannot explicitly account for persistent individual differences in reproductive success (Lee et al. 2011), nor effects of reproduction on subsequent survival that last more than one time period.

## ACKNOWLEDGEMENTS

The author is grateful to Bill Hill for many insightful discussions over the years, relating to effective population size as well as other topics. An anonymous reviewer provided comments that substantially improved the manuscript.

## Data archiving

All results presented here were generated by simulations. R code to conduct simulations for the *NegBinom* algorithm is available in Supporting Information, and all code will be deposited on Zenodo on acceptance. The author declares no conflict of interest.

## Notes

### Competing Interest Statement

The authors have declared no competing interest.

### Summary of Updates

minor changes to some results assumptions of Hill's model were clarified

