## Supplemental Info for "Robustness of Hill’s overlapping-generation method for calculating *N_e_* to extreme patterns of reproductive success"

An alternative simulation algorithm
Tables and Figures
R code
Input files

---

### An Alternative Simulation Algorithm

#### Methods

To illustrate an alternative way to generate correlations between reproduction and survival, the analyses in the main text were repeated using a different simulation algorithm. *THEWEIGHT* (Waples 2020, 2022a) is a generalized Wright-Fisher model that allows for unequal parental expectations of reproductive success, specified by a vector of parental weights, $\boldsymbol{W}$. For a given set of $b_x$, $N_x$, and $\phi_x$ values, *THEWEIGHT* randomly generates a vector of parental weights that on average will produce the desired level of reproductive skew, which depends on the coefficient of variation of the parental weights. Under Scenario LowSkew, all weights were equal so the model simplified to the standard Wright-Fisher model with Poisson variance in offspring number each time period. For Scenarios ModerateSkew and HighSkew in males, random $\boldsymbol{W}$ vectors were repeatedly generated until one was found with a coefficient of variation within a specified threshold (generally a few percent) of the target (see Waples 2022a for details). A convenient feature of *THEWEIGHT* is that it allows the user to simulate exactly the target number of offspring by randomly choosing parents for each offspring, with probabilities determined by $\boldsymbol{W}$. For each age of the adult lifespan, a pool of potential parents of the respective age was assembled, and from this pool $B_x$ parental IDs were drawn, with replacement, using the weights as relative probabilities. This process was continued until a list of 400 parental IDs had been chosen. This process was then repeated for the opposite sex, and both resulting lists were randomized before combining them to specify both parents for each offspring in the new cohort.

#### Results

In the main text, the *rnbinom* function in R was used to generate random vectors of offspring numbers. In *THEWEIGHT*, random vectors of parental weights were generated. In Model NoCor, the weights were assigned randomly to surviving individuals; otherwise, the weights were first sorted either largest-to-smallest (Model Positive) or smallest-to-largest (Model Negative) before assigning to survivors.

The two algorithms for modeling reproduction were equally successful in achieving the desired level of reproductive skew under Model NoCor, for which an analytical expectation is possible: excellent agreement was found between observed and expected $V_{k\bullet}$ for all three Scenarios (Figure 1; Table S3). As expected, under Scenario LowSkew results for *THEWEIGHT* did not vary across the three Models, whereas those for *NEGBINOM* did. For Scenarios ModerateSkew and HighSkew, reproduction based on the parental weights led to positive and negative correlations in reproductive success over time for Models Positive and Negative, respectively, but with slightly different magnitudes than obtained with *NegBinom* (compare Table 3 for *NegBinom* and Table S2 for *TheWeight*).

Two factors contributed to these relatively modest differences. First, whereas in *NegBinom* the vectors of randomly-generated offspring numbers were sorted before assigning to individuals, in *TheWeight* what was sorted were randomly-generated parental weights, which only represent relative probabilities of producing offspring, not actual offspring numbers. The second factor is that when reproductive skew is very high ($\phi$=20 in Scenario HighSkew), most randomly-generated weights, and most randomly-generated offspring numbers, will be 0. As discussed by Waples (2022a), this creates some challenges when modeling age-structured populations of long-lived species, for which the numbers of individuals in older age classes can be very small (in the core lifetable in Table 1, there are only 8 age-10 individuals of each sex).

Results show that either algorithm can be used effectively to generate covariance between reproduction and survival and/or temporal autocorrelations in reproductive success. Here is a quick summary of some of the pros and cons of the two algorithms:

*NegBinom*

Pros

- Fast; only requires generating a vector of random offspring numbers, given a specified mean and variance.
- Numerous examples of this general approach are in the published literature.

Cons

- Total number of offspring produced is a random variable, which might require some adjustments depending on the experimental design.
- Realized offspring numbers (fitnesses) are randomly assigned to individuals rather than reflecting any individual traits.

*TheWeight*

Pros

- Total number of offspring produced is specified by user.
- Parental weights are a natural way to map relative fitnesses to specific individuals based on individual trait values.

Cons

- Slower; before generating offspring, must randomly generate parental weights with the desired properties.
- Recently proposed; only a few examples of this general approach are in the published literature.

Table S1. As in Table 2, but showing results for Model Negative. The left set of columns replicate those in Table 2, where offspring numbers were randomly generated for survivors at each age. In the right set of columns, those random numbers were sorted so that each year the individual with the highest ID produced the most offspring. As individuals with the highest IDs died before reaching the next age, this created negative correlations between reproduction and survival. The columns labeled 'LRS' show the total lifetime numbers of offspring produced by each individual, and the columns labeled 'LRS-' show LRS for all ages except age 1. See text for discussion.

| ID | 1 | 2 | 3 | 4 | 5 | LRS | LRS- | 1 | 2 | 3 | 4 | 5 | LRS | LRS- |
|---|---|---|---|---|---|---|---|---|---|---|---|---|---|---|
| 1 | 6 | 5 | 1 | 1 | 4 | 17 | 11 | 0 | 0 | 0 | 1 | 4 | 5 | 5 |
| 2 | 0 | 1 | 0 | 6 | 4 | 11 | 11 | 0 | 0 | 0 | 1 | 4 | 5 | 5 |
| 3 | 0 | 1 | 2 | 1 | NA | 4 | 4 | 0 | 0 | 0 | 5 | NA | 5 | 5 |
| 4 | 1 | 0 | 0 | 8 | NA | 9 | 8 | 0 | 0 | 1 | 6 | NA | 7 | 7 |
| 5 | 1 | 17 | 8 | 5 | NA | 31 | 30 | 0 | 1 | 2 | 8 | NA | 11 | 11 |
| 6 | 0 | 0 | 0 | NA | NA | 0 | 0 | 0 | 1 | 3 | NA | NA | 4 | 4 |
| 7 | 0 | 0 | 7 | NA | NA | 7 | 7 | 0 | 1 | 7 | NA | NA | 8 | 8 |
| 8 | 0 | 0 | 3 | NA | NA | 3 | 3 | 0 | 1 | 8 | NA | NA | 9 | 9 |
| 9 | 0 | 1 | NA | NA | NA | 1 | 1 | 0 | 1 | NA | NA | NA | 1 | 1 |
| 10 | 0 | 2 | NA | NA | NA | 2 | 2 | 0 | 2 | NA | NA | NA | 2 | 2 |
| 11 | 2 | 1 | NA | NA | NA | 3 | 1 | 0 | 5 | NA | NA | NA | 5 | 5 |
| 12 | 0 | 1 | NA | NA | NA | 1 | 1 | 0 | 17 | NA | NA | NA | 17 | 17 |
| 13 | 0 | NA | NA | NA | NA | 0 | 0 | 0 | NA | NA | NA | NA | 0 | 0 |
| 14 | 0 | NA | NA | NA | NA | 0 | 0 | 0 | NA | NA | NA | NA | 0 | 0 |
| 15 | 5 | NA | NA | NA | NA | 5 | 0 | 1 | NA | NA | NA | NA | 1 | 0 |
| 16 | 0 | NA | NA | NA | NA | 0 | 0 | 1 | NA | NA | NA | NA | 1 | 0 |
| 17 | 3 | NA | NA | NA | NA | 3 | 0 | 2 | NA | NA | NA | NA | 2 | 0 |
| 18 | 0 | NA | NA | NA | NA | 0 | 0 | 3 | NA | NA | NA | NA | 3 | 0 |
| 19 | 0 | NA | NA | NA | NA | 0 | 0 | 5 | NA | NA | NA | NA | 5 | 0 |
| 20 | 0 | NA | NA | NA | NA | 0 | 0 | 6 | NA | NA | NA | NA | 6 | 0 |

Table S2. As in Table 3, main text, but showing simulation results for *THEWEIGHT* algorithm. Columns on the left show the $N_e$ calculated from the simulated pedigrees; middle columns show the ratio of observed heterozygosity to expected heterozygosity based on the pedigree $N_e$; columns on the right show the correlation ($\rho_{\alpha,\alpha+}$) between offspring numbers for each individual at age α and for all subsequent ages combined (as shown in the *LRS*- column from Table 2). Pedigree $N_e$ was calculated as the harmonic mean across cohorts and replicates; the ratio $H_{obs}/H_{exp}$ was the average across 10 replicates, and within each replicate the ratio was calculated as the mean across the final 50 years. For each cohort in each replicate, $\rho_{\alpha,\alpha+}$ was computed across all individuals that reached age at maturity, and results were averaged across cohorts and replicates.

| | | | Pedigree $N_e$ | | | $H_{obs}/H_{exp}$ | | | Correlation ($\rho_{\alpha,\alpha+}$) | | | |
|---|---|---|---|---|---|---|---|---|---|---|---|---|
| | Model NoCor | | Model | | | Model | | | Model Positive | | Model Negative | |
| Scenario | E($V_{k\bullet}$) | E($N_e$) | NoCor | Positive | Negative | NoCor | Positive | Negative | F | M | F | M |
| LowSkew | 10.2 | 636 | 640 | 640 | 639 | 1.001 | 1.001 | 0.999 | 0.001 | -0.006 | -0.006 | 0.004 |
| ModerateSkew | 16.1 | 462 | 468 | 201 | 673 | 1.007 | 0.994 | 1.003 | | 0.91 | | -0.37 |
| HighSkew | 31.1 | 253 | 263 | 65 | 329 | 0.998 | 1.021 | 0.999 | | 0.90 | | -0.09 |

Table S3. Mean values of the lifetime variance in reproductive success ($V_{k\bullet}$) for simulated data using two different simulation algorithms. Observed and expected results are shown for Model NoCor, for which analytical results are possible based on the vital rates in Table 1. The expected and observed *NEGBINOM* results are illustrated in Figure 1.

| | LowSkew | ModerateSkew | HighSkew |
|---|---|---|---|
| Observed - *NEGBINOM* | 10.17 | 21.94 | 51.37 |
| Observed - *THEWEIGHT* | 10.17 | 21.76 | 49.71 |
| Expected | 10.18 | 21.95 | 51.95 |

Table S4. The ratio of observed to expected $\Delta V_{P(t)}$ in simulations using the negative binomial algorithm. $\Delta V_{P(t)}$ is the increase in allele frequency variance $t$ generations after the burnin period (which included years 1-50). The ratio is averaged across years 451-500 in 10 replicates. See Table 3 for comparable results for the rate of loss of heterozygosity. The average ratio across models and scenarios is 1.007.

| | | Model | |
|---|---|---|---|
| **Scenario** | **NoCor** | **Positive** | **Negative** |
| LowSkew | 0.975 | 1.002 | 1.056 |
| ModerateSkew | 0.977 | 0.985 | 1.029 |
| HighSkew | 1.003 | 0.994 | 1.042 |

Table S5. Vital rates for the 5- and 20-year alternative-lifetable scenarios whose results are shown in Figure S1.

| | Female | | | Male | | | Female | | | Male | | |
|---|---|---|---|---|---|---|---|---|---|---|---|---|
| Age | sx | bx | phi | sx | bx | phi | sx | bx | phi | sx | bx | phi |
| 1 | 0.85 | 0 | 1 | 0.85 | 0 | 5 | 0.5 | 1 | 1 | 0.5 | 1 | 5 |
| 2 | 0.85 | 0 | 1 | 0.85 | 0 | 5 | 0.5 | 1 | 1 | 0.5 | 2 | 5 |
| 3 | 0.85 | 0 | 1 | 0.85 | 0 | 5 | 0.5 | 1 | 1 | 0.5 | 3 | 5 |
| 4 | 0.85 | 0 | 1 | 0.85 | 0 | 5 | 0.5 | 1 | 1 | 0.5 | 4 | 5 |
| 5 | 0.85 | 1 | 1 | 0.85 | 5 | 5 | 0.5 | 1 | 1 | 0.5 | 5 | 5 |
| 6 | 0.85 | 1 | 1 | 0.85 | 6 | 5 | | | | | | |
| 7 | 0.85 | 1 | 1 | 0.85 | 7 | 5 | | | | | | |
| 8 | 0.85 | 1 | 1 | 0.85 | 8 | 5 | | | | | | |
| 9 | 0.85 | 1 | 1 | 0.85 | 9 | 5 | | | | | | |
| 10 | 0.85 | 1 | 1 | 0.85 | 10 | 5 | | | | | | |
| 11 | 0.85 | 1 | 1 | 0.85 | 11 | 5 | | | | | | |
| 12 | 0.85 | 1 | 1 | 0.85 | 12 | 5 | | | | | | |
| 13 | 0.85 | 1 | 1 | 0.85 | 13 | 5 | | | | | | |
| 14 | 0.85 | 1 | 1 | 0.85 | 14 | 5 | | | | | | |
| 15 | 0.85 | 1 | 1 | 0.85 | 15 | 5 | | | | | | |
| 16 | 0.85 | 1 | 1 | 0.85 | 16 | 5 | | | | | | |
| 17 | 0.85 | 1 | 1 | 0.85 | 17 | 5 | | | | | | |
| 18 | 0.85 | 1 | 1 | 0.85 | 18 | 5 | | | | | | |
| 19 | 0.85 | 1 | 1 | 0.85 | 19 | 5 | | | | | | |
| 20 | 0.85 | 1 | 1 | 0.85 | 20 | 5 | | | | | | |

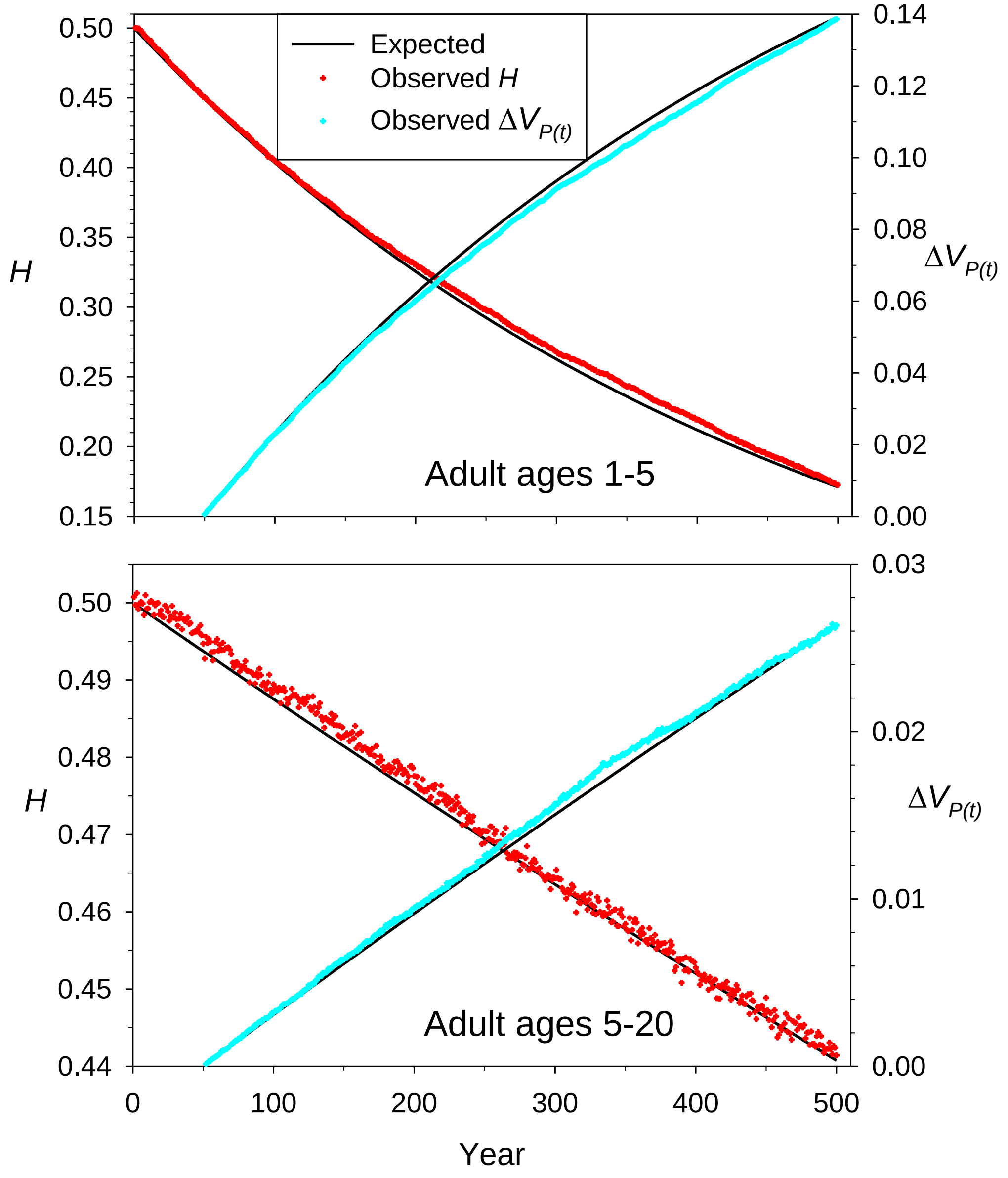


Figure S1. Observed (colored symbols) and expected (black lines) rates of genetic drift in simulated populations using either a 5-year lifespan with α=1 (top panel) or 20-year lifespan with α=5 (bottom panel) (see Table S5 for details of the vital rates). These simulations used Scenario ModerateSkew under Model Positive, and harmonic mean pedigree $N_e$ was 108.8 (top) and 194.8 (bottom) .

### R code to implement the NegBinom simulation algorithm

```
### This code uses negative binomial function for reproduction
### For each of NReps replicates, NYears of annual reproduction is modeled.
### The code keeps track of demography and tracks genetic variation at NLoci diallelic ('SNP') loci
## Note that this code was modified from code that used parental weights, so dummy weights are generated but
not used

library(plyr) ## required for the 'count' function

######## User Inputs
Vitals = read.csv(file="inputfilename")
Vitals = read.csv(file="C:/users/lukes/dropbox/HillTests/TestingHill5.csv")
Method = "NegBin"
N1 = 200 # number of yearlings of each sex in each cohort
NLoci = 100
NYears = 500
SimNe = c(635.8,462.1,252.5)
HillNe =SimNe[2]  ## from AgeNe, based on input data
Option = 2 #### 1 = NoCor model; 2 = positive autocorrelations and covariances; 3 = negative autocorrelations and
covariances
NReps =2

###############
demography <- function(omega,Survival,N1,Femalebx,Malebx)  {
 S = c(1,Survival[-omega])
 Lx = cumprod(S)       ### cumulative survival
 Nx = round(N1*Lx,0)   ### expected number in each age class
 FIndex = sum(Femalebx*Lx)  ## should be 2 in a stable pop with equal sex ratio
 Femalebx2 = 2*Femalebx/FIndex
 MIndex = sum(Malebx*Lx)  ## should be 2 in a stable pop with equal sex ratio
 Malebx2 = 2*Malebx/MIndex
  D = cbind(Lx,Nx,Femalebx2,Malebx2)
 return(D)   }  ## end function

################
initialize <- function(alpha,omega,NLoci,Nx,Initialmu)   {
  NTot = sum(Nx)
  Adults = Nx[alpha:omega]
  AL = omega-alpha+1
  G = matrix(1,NTot,NLoci)
    Age = 9999
  for (j in 1:omega) {  ## assign ages
    Age = c(Age,rep(j,Nx[j]))
    }  # end for j
  Age = Age[-1]
 ID = rev(1:NTot) ## assign unique ID
 ## Get initial weights
 W = rep(0,NTot)
  end = cbind(ID,Age,W,G)  ## metadata for each individual
 return(end)   }  ## end function
```

```
#######
get_parents <- function(MaleBit,FemaleBit,Malebxprime,Femalebxprime) {
AgeData = matrix(NA,AL,4)
upper = 1.05*cohortsize
lower = cohortsize

repeat {
   Dads = 9999
  for (j in 1:AL)  { ## pick IDs for dads
    someDads = subset(MaleBit,MaleBit[,"Age"] == AdultAges[j])
    someIDs = someDads[,"ID"]
    MVk = Malephi[j]*Malebxprime[j]
  if (Malephi[j] == 1) {kvector <- rpois(AdultSurvivors[j], Malebxprime[j])}  ## use Poisson distribution for phi = 1 and
negative binomial for phi > 1
    else { kvector <- rnbinom(AdultSurvivors[j], mu = Malebxprime[j], size = Malebxprime[j]^2/(MVk-
Malebxprime[j])) }
  AgeData[j,1] = mean(kvector)
  AgeData[j,2] = var(kvector)
  if (Option == 2)  { kvector = rev(sort(kvector))  }
  else if (Option == 3)  { kvector = sort(kvector)  }
        for (k in 1:length(kvector)) {
          add = rep(someDads[k],kvector[k])
          Dads = c(Dads,add)
          }  # end for k
    }  # end for j
    Dads = Dads[-1]
  if(length(Dads) > lower & length(Dads) < upper)  { break}
  }  # end repeat
    Dads = sample(Dads,cohortsize,replace=F) ##  trim list of dads to desired cohortsize and randomize order

  ## repeat for Moms
  repeat {
    Moms = 9999
   for (j in 1:AL)  {
      someMoms = subset(FemaleBit,FemaleBit[,"Age"] == AdultAges[j])
      someIDs = someMoms[,"ID"]
      FVk = Femalephi[j]*Femalebxprime[j]
    if (Femalephi[j] == 1) {kvector <- rpois(AdultSurvivors[j], Femalebxprime[j])}  ## use Poisson distribution for phi =
1 and negative binomial for phi > 1
      else { kvector <- rnbinom(AdultSurvivors[j], mu = Femalebxprime[j], size = Femalebxprime[j]^2/(FVk-
Femalebxprime[j])) }
    AgeData[j,3] = mean(kvector)
    AgeData[j,4] = var(kvector)
    if (Option == 2)  { kvector = rev(sort(kvector))  }
    else if (Option == 3)  { kvector = sort(kvector)  }
      for (k in 1:length(kvector)) {
         add = rep(someMoms[k],kvector[k])
         Moms = c(Moms,add)
         }  # end for k
    }  # end for j
    Moms = Moms[-1]
     if(length(Moms) > lower & length(Moms) < upper)  { break}
  }  # end repeat
```

```
    Moms = sample(Moms,cohortsize,replace=F)

DadRows = rep(NA,cohortsize)
MomRows = DadRows
   for (i in 1:cohortsize)  {
    DadRows[i] = which(MaleBit[,"ID"] == Dads[i])  ## row number for each parent
    MomRows[i] = which(FemaleBit[,"ID"] == Moms[i])
    }  ## end for i
  BigParents = rbind(DadRows,MomRows)

  MyList <- list("a" = BigParents, "b" = AgeData)
  return(MyList)    } ## end function

############
reproduce <- function(MaleGenosPlus,FemaleGenosPlus,BigParents,Y){
  MaleGenos = MaleGenosPlus[,-1:-3]
  FemaleGenos = FemaleGenosPlus[,-1:-3]
  Malebit = MaleGenosPlus[,1:3]
  Femalebit = FemaleGenosPlus[,1:3]

  NOffspring = ncol(BigParents)
  L = dim(MaleGenos)[[2]] # number of loci (columns)
  Dads = BigParents[1,]
  Moms = BigParents[2,]

  # construct matrices of the parental genotypes and metadata
  parents_geno1 = MaleGenos[Dads, ]
  parents_geno2 = FemaleGenos[Moms, ]
  DadMeta = Malebit[Dads,]
  DBY = Y-DadMeta[,"Age"]
  DadMeta = cbind(DadMeta[,1],DadMeta[,2],DBY,DadMeta[,3])
  MomMeta = Femalebit[Moms,]
  MBY = Y - MomMeta[,"Age"]
  MomMeta = cbind(MomMeta[,1],MomMeta[,2],MBY,MomMeta[,3])
  BigMeta = cbind(DadMeta,MomMeta)
  colnames(BigMeta) = c("DadID","DadAge","DadBirthYear", "DadW","MomID","MomAge","MomBirthYear",
"MomW")

  # offspring are a combination of the two parents
    offspring <- rbinom(n=NOffspring*NLoci, size = 1, p = parents_geno1 / 2) +
      rbinom(n = NOffspring*NLoci, size = 1, p = parents_geno2 / 2)
    # convert offspring vector back to a matrix
  offspring <- matrix(data = offspring, nrow =NOffspring, ncol = L )

  offspringplus = cbind(BigMeta,offspring)

  return(offspringplus)
         } # end function

######################
mortality <- function(Genos,survival,Nx)   {
  survivors = rep(9999,ncol(Genos))
  temp = subset(Genos, Genos[,"Age"] < omega)  ### all individual in oldest age class die
```

```
  for (j in 1:(omega-1))  {
    X = subset(temp,temp[,"Age"] == j)
    Y = X[1:Nx[j+1],]  ### first individuals in each age class survive
    survivors = rbind(survivors,Y)
    }  ## end for j
  survivors = survivors[-1,]
  survivors[,"Age"] = survivors[,"Age"] + 1  ## survivors all age 1 year
return(survivors)  }  ## end function

######################

#########  Main Program

omega = nrow(Vitals)  # maximum age
AL = sum(Vitals$Femalebx>0) ## adult lifespan; same as for males in this example
alpha = omega - AL + 1  # fixed age at first reproduction
Ages = 1:omega
AdultAges = Ages[alpha:omega]
cohortsize = 2*N1
Sex = c(1,2)  ## 1 = Male; 2 = Female
Gender = c(rep(Sex[1],N1),rep(Sex[2],N1)) ## fixed 1:1 sex ratio

Malebx = Vitals$Malebx
Malephi = Vitals$Malephi[alpha:omega]
Femalebx = Vitals$Femalebx
Femalephi = Vitals$Femalephi[alpha:omega]
survival = Vitals$FemaleSx ## same as for males in this example

demo = demography(omega,survival,N1,Femalebx,Malebx)
lx = demo[,1]
NSurvivors = demo[,2]
AdultSurvivors = NSurvivors[alpha:omega]
AdultN = sum(AdultSurvivors)
TotN = sum(NSurvivors)
Femalebxprime = demo[alpha:omega,3] ## fecundity rescaled to produce constant N
Malebxprime = demo[alpha:omega,4]
FemaleBx = round(AdultSurvivors*Femalebxprime)
MaleBx = round(AdultSurvivors*Malebxprime)
FemaleBx[1] = cohortsize - sum(FemaleBx[2:AL])
MaleBx[1] = cohortsize - sum(MaleBx[2:AL])

HugeH = matrix(NA,NYears,NReps)
HugeP = array(NA,dim = c(NYears,NLoci,NReps))

HugeVk = matrix(NA,NReps,2)
colnames(HugeVk) = c("Female","Male")
HugePhi = array(NA, dim = c(NReps,AL,2))
colnames(HugePhi) = AdultAges
dimnames(HugePhi)[[3]] = c("Female","Male")

for (R in 1:NReps)  {

H = rep(NA,NYears)  ## tracks mean heterozygosity
```

```
P = matrix(NA,NYears,NLoci)
Pedigree = matrix(9999,1,12)
colnames(Pedigree) = c("Year", "YearlingID", "gender", "YearlingW", "DadID", "DadAge", "DadBirthYear", "DadW",
"MomID", "MomAge", "MomBirthYear", "MomW")

BigAgeData = array(NA, dim = c(AL,4,NYears))
colnames(BigAgeData) = c("Male bx","Male Vk","Female bx","Female Vk")
rownames(BigAgeData) = AdultAges

report = (NYears/10)*(1:10)  ### report progress every 10% of job

#### Initialize population
MaleGenosPlus = initialize(alpha,omega,NLoci,NSurvivors,Malemu)
FemaleGenosPlus = initialize(alpha,omega,NLoci,NSurvivors,Femalemu)


for (Y in 1:NYears)  {

if(Y %in% report)  {
   print(paste0("Rep = ",R,"  Year = ",Y))
   flush.console()    }  ## end if

AdultMales = subset(MaleGenosPlus,MaleGenosPlus[,"Age"]>=alpha)
AdultFemales = subset(FemaleGenosPlus,FemaleGenosPlus[,"Age"]>=alpha)

ParentalStuff <- get_parents(AdultMales[,1:3],AdultFemales[,1:3],Malebxprime,Femalebxprime)
 BothParents = ParentalStuff[[1]]
 NewAgeData = ParentalStuff[[2]]
 BigAgeData[,,Y] = NewAgeData

cohort = reproduce(AdultMales,AdultFemales,BothParents,Y)

newgenos = cohort[,-1:-8]
H[Y] = table(newgenos)[2]/(nrow(newgenos)*ncol(newgenos))  ## mean observed heterozygosity
P[Y,] = colMeans(newgenos)/2 ## allele frequencies
newped = cohort[,1:8]
newmeta = matrix(NA,nrow(cohort),3)
Gender = sample(Gender,nrow(cohort),replace=FALSE) ## randomly pick gender of each offspring
Year = rep(Y,nrow(cohort))
YearlingID = rep(NA,nrow(cohort))
YearlingW = YearlingID
newPlus = matrix(NA,nrow(cohort),3)
colnames(newPlus) = c("ID","Age","W")
Big = cbind(Year,YearlingID,Gender,YearlingW,newped,newPlus,newgenos)
newMales = subset(Big,Big[,"Gender"] == 1)
newFemales = subset(Big,Big[,"Gender"] == 2)

femW = rep(0,nrow(newFemales))
newFemales[,"YearlingW"] = femW
newFemales[,"W"] = femW
maleW = rep(0,nrow(newMales))
newMales[,"YearlingW"] = maleW
newMales[,"W"] = maleW
```

```
NextID = TotN + round((Y-1)*1.05*cohortsize)
newMaleIDs = (NextID+1):(NextID+nrow(newMales))
newMales[,"YearlingID"] = newMaleIDs
newMales[,"ID"] = newMaleIDs
newMales[,"Age"] = rep(1,nrow(newMales))
Pedigree = rbind(Pedigree,newMales[,1:12])
newMaleGenos = newMales[,-1:-12]

newFemaleIDs = (NextID+1):(NextID+nrow(newFemales))
newFemales[,"YearlingID"] = newFemaleIDs
newFemales[,"ID"] = newFemaleIDs
newFemales[,"Age"] = rep(1,nrow(newFemales))
Pedigree = rbind(Pedigree,newFemales[,1:12])
newFemaleGenos = newFemales[,-1:-12]

survivingmales <- mortality(MaleGenosPlus,survival,NSurvivors)
MaleGenosPlus = rbind(newMaleGenos,survivingmales)

survivingfemales <- mortality(FemaleGenosPlus,survival,NSurvivors)
FemaleGenosPlus = rbind(newFemaleGenos,survivingfemales)

}  # end for Y

Pedigree = Pedigree[-1,]

HugeH[,R] = H
HugeP[,,R] = P

## can write the Pedigree file to disk for later analysis

############################################## Plot loss of H over time, compared to rate expected based on Ne calculated in AgeNe
BigGen = 5.218

ObsVitals = matrix(NA,AL,6)  #3 mean age-specific vital rates in simulations
colnames(ObsVitals) = c("Male bx","Male Vk","Female bx","Female Vk","Male phi","Female phi")
rownames(ObsVitals) = AdultAges
for (j in 1:AL)  {
  for (k in 1:4)  { ObsVitals[j,k] = mean(BigAgeData[j,k,]) }
  ObsVitals[j,5] = ObsVitals[j,2]/ObsVitals[j,1]
  ObsVitals[j,6] = ObsVitals[j,4]/ObsVitals[j,3]
  } # end for j

HugePhi[R,,1] = ObsVitals[,"Female phi"]
HugePhi[R,,2] = ObsVitals[,"Male phi"]

MaleNx = NSurvivors
FemaleNx = NSurvivors
AdultMaleNx = AdultSurvivors
AdultFemaleNx = AdultSurvivors

##lifetime reprodutive success
```

```
## exclude first burnin and last years
Start = 1 + 2*omega
End = NYears - 2*omega
tot = End-Start + 1
Malekbar = seq(1:tot)
Femalekbar = seq(1:tot)
MaleVk = seq(1:tot)
FemaleVk = seq(1:tot)
OffspringM = seq(1:tot)
OffspringF = seq(1:tot)
MaleGen = seq(1:tot)
FemaleGen = seq(1:tot)
MaxStud = seq(1:tot)
MaxMama = seq(1:tot)
Studs = 1:tot
Mamas = 1:tot

MaleCohort = MaleNx[1]
FemaleCohort = MaleCohort

MaleCor = 1:tot
FemaleCor = 1:tot
current = Start
for (i in 1:tot) {
MaleLRO = matrix(0,MaleCohort,2)
FemaleLRO = matrix(0,FemaleCohort,2)
MV <- subset(Pedigree, Pedigree[,"DadBirthYear"] == current)
MV1 <- count(MV[,"DadID"])
  for(j in 1:nrow(MV1))  {
   A = subset(MV,MV[,"DadID"]== MV1$x[j])  ### LRO for that dad
   B = subset(A,A[,"DadAge"]==alpha)  ### first reproduction for that dad
   MaleLRO[j,1] = nrow(B)
   MaleLRO[j,2] = nrow(A) - nrow(B)
    }  ## end for j
MaxStud[i] <- max(MV1$freq)
OffspringM[i] <- sum(MV1$freq)  ## total number of lifetime offspring produced by the cohort of males
Malekbar[i] <- sum(MV1$freq)/MaleCohort
Studs[i] <- nrow(MV1)  ## number of males in cohort that ever produced offspring
zeros <- MaleCohort - Studs[i]  ## males in the cohort that were duds for a lifetime
MV2 <- MV1$freq^2
sum(MV2)
MaleVk[i] <- sum(MV2)/MaleCohort - Malekbar[i]^2
MaleGen[i] <- mean(MV[,"DadAge"])

FV <- subset(Pedigree, Pedigree[,"MomBirthYear"] == current)
FV1 <- count(FV[,"MomID"])
 for(j in 1:nrow(FV1))  {
   AA = subset(FV,FV[,"MomID"]== FV1$x[j])
   BB = subset(AA,AA[,"MomAge"]==alpha)
   FemaleLRO[j,1] = nrow(BB)
   FemaleLRO[j,2] = nrow(AA) - nrow(BB)
   }  ## end for j
MaxMama[i] <- max(FV1$freq)
```

```
OffspringF[i] <- sum(FV1$freq)  ## total number of lifetime offspring produced by the cohort of females
Femalekbar[i] <- sum(FV1$freq)/FemaleCohort
Mamas[i] <- nrow(FV1)  ## number of females in cohort that ever produced offspring
zeros <- FemaleCohort - Mamas[i]  ## females in the cohort that were duds for a lifetime
FV2 <- FV1$freq^2
sum(FV2)  ## sum(ki^2) from PWOP
FemaleVk[i] <- sum(FV2)/FemaleCohort - Femalekbar[i]^2
FemaleGen[i] <- mean(FV[,"MomAge"])
current <- current + 1

MaleCor[i] = cor(MaleLRO[,1],MaleLRO[,2])
FemaleCor[i] = cor(FemaleLRO[,1],FemaleLRO[,2])
} # end for i

##Rescale Vk to expected value in stable population
MaleVk2 = 2*(1 + 2*(MaleVk/Malekbar - 1)/Malekbar)
FemaleVk2 = 2*(1 + 2*(FemaleVk/Femalekbar - 1)/Femalekbar)
MaleNe = 4*MaleCohort*MaleGen/(2 + MaleVk2)
FemaleNe = 4*FemaleCohort*FemaleGen/(2 + FemaleVk2)

z <- cbind(Start:End,MaleCohort, Studs,OffspringM, MaxStud, Malekbar, MaleVk, MaleGen, MaleVk2,MaleNe)
zz <- cbind(Start:End, FemaleCohort, Mamas, OffspringF, MaxMama, Femalekbar, FemaleVk, FemaleGen,
FemaleVk2,FemaleNe)
colnames(z) <- c("Year","Males", "Sires", "NOffspring", "BigStud", "kbar", "Vk", "Gen", "Vk2","Ne")
colnames(zz) <- c("Year", "Females", "Mamas", "NOffspring", "MaxMama", "kbar", "Vk", "Gen", "Vk2","Ne")

#head(z)
#head(zz)

BigGen = (mean(MaleGen)+mean(FemaleGen))/2
BigVk2 = (mean(MaleVk2)+ mean(FemaleVk2))/2
BigMaleNe = 1/mean(1/MaleNe) ## harmonic mean across cohorts
BigFemaleNe = 1/mean(1/FemaleNe)
BigNe = 4*cohortsize*BigGen/(2+BigVk2)

Summary = matrix(NA,4,3)
rownames(Summary) = c("Nb","Generation","Lifetime Vk*","Ne")
colnames(Summary) = c("Female","Male","Overall")

Summary[2,] = c(mean(FemaleGen), mean(MaleGen), BigGen)
Summary[3,] = c(mean(FemaleVk2), mean(MaleVk2), BigVk2)
Summary[4,] = c(BigFemaleNe,BigMaleNe,BigNe)

HugeVk[R,1] = mean(FemaleVk2)
HugeVk[R,2] = mean(MaleVk2)

##Observed demographic results in simulations
#ObsVitals
#colMeans(z)
#colMeans(zz)
#Summary

##PedigreeNe = Summary[4,3]
```

```
#mean(MaleCor)
#mean(FemaleCor)
#c(HillNe,PedigreeNe)

}  # end for R

OverallH = rowMeans(HugeH)
OverallVk = (mean(HugeVk[,1])+mean(HugeVk[,2]))/2
PedigreeNe = 4*2*N1*BigGen/(2+OverallVk)

ExpH1 = 1:NYears
ExpH2 = ExpH1
for (j in 1:NYears)  {
  t = j/BigGen
  ExpH1[j] = 0.5*(1 - 1/(2*HillNe))^t
  ExpH2[j] = 0.5*(1 - 1/(2*PedigreeNe))^t
   }

Varp = matrix(NA,NYears,NReps)

for(Y in 50:NYears)  {
for(R in 1:NReps)  {
  Varp[Y,R] = var(HugeP[Y,,R])
  }}
VarpBar = rowMeans(Varp)
BurnVarp = VarpBar[50]
DiffVarp = VarpBar-BurnVarp

PZero = HugeP[50,,]
PP = PZero*(1-PZero)
PPbar = mean(PP)

EDiffVarp = rep(NA,NYears)
for (j in 51:NYears)  {
  t = (j-50)/BigGen
EDiffVarp[j] = PPbar*(1 - (1 - 1/(2*PedigreeNe))^t)
}

ExpH1 = 1:NYears
ExpH2 = ExpH1
for (j in 1:NYears)  {
  t = j/BigGen
  ExpH1[j] = 0.5*(1 - 1/(2*HillNe))^t
  ExpH2[j] = 0.5*(1 - 1/(2*PedigreeNe))^t
   }

colMeans(HugePhi)
c(HillNe,PedigreeNe,PedigreeNe/HillNe)
Hetsbit = cbind(OverallH[(NYears-49):NYears],ExpH2[(NYears-49):NYears])
mean(Hetsbit[,1]/Hetsbit[,2])
Varbit = cbind(DiffVarp[(NYears-49):NYears],EDiffVarp[(NYears-49):NYears])
mean(Varbit[,1]/Varbit[,2])
```

```
#cbind(DiffVarp,EDiffVarp)
#cbind(EDiffVarp,DiffVarp)

##plot actual vs expected H and Var(P)

bottom = min(OverallH)*0.99
### plot actual loss of heterozygosity (red triangles) and predicted loss (dashed blue line) over time
plot(1:NYears,OverallH,type="p",lwd=3,col="red",xlab="Year",ylim = c(bottom,0.505),ylab="Heterozygosity")
lines(1:NYears,ExpH1,type="l",lty=3,lwd=3,col="blue")
lines(1:NYears,ExpH2,type="l",lty=3,lwd=3,col="green")
legend(0.48,0.48, legend=c("Actual", "Predicted NoCor","Pedigree"),col=c("red", "blue","green"), lty=1:2, cex=1)

top = max(DiffVarp,na.rm=T)*1.5
### plot actual var(P) (red triangles) and predicted loss (dashed blue line) over time
plot(1:NYears,DiffVarp,type="p",lwd=3,col="cyan",xlab="Year",ylim = c(0,top),ylab="Delta(Vp)")
lines(1:NYears,EDiffVarp,type="l",lty=3,lwd=3,col="blue")
legend(0.1,0.05, legend=c("Actual", "Expected"),col=c("cyan", "blue"), lty=1:2, cex=1)
```

### Input files for simulations

#### Scenario LowSkew

| Age | FemaleSx | Femalebx | Femalephi | MaleSx | Malebx | Malephi |
|---|---|---|---|---|---|---|
| 1 | 0.7 | 0 | 0 | 0.7 | 0 | 0 |
| 2 | 0.7 | 0 | 0 | 0.7 | 0 | 0 |
| 3 | 0.7 | 1 | 1 | 0.7 | 1 | 1 |
| 4 | 0.7 | 1 | 1 | 0.7 | 1 | 1 |
| 5 | 0.7 | 1 | 1 | 0.7 | 1 | 1 |
| 6 | 0.7 | 1 | 1 | 0.7 | 1 | 1 |
| 7 | 0.7 | 1 | 1 | 0.7 | 1 | 1 |
| 8 | 0.7 | 1 | 1 | 0.7 | 1 | 1 |
| 9 | 0.7 | 1 | 1 | 0.7 | 1 | 1 |
| 10 | 0 | 1 | 1 | 0 | 1 | 1 |

#### Scenario ModerateSkew

| Age | FemaleSx | Femalebx | Femalephi | MaleSx | Malebx | Malephi |
|---|---|---|---|---|---|---|
| 1 | 0.7 | 0 | 0 | 0.7 | 0 | 0 |
| 2 | 0.7 | 0 | 0 | 0.7 | 0 | 0 |
| 3 | 0.7 | 1 | 1 | 0.7 | 3 | 5 |
| 4 | 0.7 | 1 | 1 | 0.7 | 4 | 5 |
| 5 | 0.7 | 1 | 1 | 0.7 | 5 | 5 |
| 6 | 0.7 | 1 | 1 | 0.7 | 6 | 5 |
| 7 | 0.7 | 1 | 1 | 0.7 | 7 | 5 |
| 8 | 0.7 | 1 | 1 | 0.7 | 8 | 5 |
| 9 | 0.7 | 1 | 1 | 0.7 | 9 | 5 |
| 10 | 0 | 1 | 1 | 0 | 10 | 5 |

#### Scenario HighSkew

| Age | FemaleSx | Femalebx | Femalephi | MaleSx | Malebx | Malephi |
|---|---|---|---|---|---|---|
| 1 | 0.7 | 0 | 0 | 0.7 | 0 | 0 |
| 2 | 0.7 | 0 | 0 | 0.7 | 0 | 0 |
| 3 | 0.7 | 1 | 1 | 0.7 | 3 | 20 |
| 4 | 0.7 | 1 | 1 | 0.7 | 4 | 20 |
| 5 | 0.7 | 1 | 1 | 0.7 | 5 | 20 |
| 6 | 0.7 | 1 | 1 | 0.7 | 6 | 20 |
| 7 | 0.7 | 1 | 1 | 0.7 | 7 | 20 |
| 8 | 0.7 | 1 | 1 | 0.7 | 8 | 20 |
| 9 | 0.7 | 1 | 1 | 0.7 | 9 | 20 |
| 10 | 0 | 1 | 1 | 0 | 10 | 20 |
